# CryoFSL: An Annotation-Efficient, Few-Shot Learning Framework for Robust Protein Particle Picking in Cryo-EM Micrographs

**DOI:** 10.1101/2025.09.19.677446

**Authors:** Biplab Poudel, Rajan Gyawali, Ashwin Dhakal, Jianlin Cheng, Dong Xu

## Abstract

Accurate identification of protein particles in cryo-electron microscopy (cryo-EM) micrographs is crucial for high-resolution structure determination, but remains challenging due to the heavy reliance on extensive annotated datasets and the difficulty of ensuring robustness under low signal-to-noise ratio (SNR) conditions. Current approaches require large annotations and exhibit poor generalization to new protein targets. We present CryoFSL (<u>Cryo</u>-EM <u>F</u>ew <u>S</u>hot-<u>L</u>earning), a novel few-shot learning framework built upon Segment Anything Model 2 (SAM2) with lightweight adapters, enabling robust particle picking using as few as five labeled micrographs, significantly reducing annotation burden. The framework’s hierarchical adapter design supports dynamic feature modulation for low-SNR and heterogeneous conditions, resolving the trade-off between annotation burden and performance. CryoFSL surpasses both traditional template-based methods and state-of-the-art deep learning models across diverse proteins in the few-short learning setting, achieving superior recall, precision and 3D reconstruction resolution with minimal supervision. It maintains stability across heterogeneous micrographs and consistently detects high-quality particles with fewer false positives. Notably, CryoFSL achieves competitive density map reconstruction resolution with just a fraction of the particles picked by other methods, redefining efficiency and quality in cryo-EM analysis. This work paves the way for scalable, generalizable, and annotation-efficient particle picking pipelines. The code is available at GitHub.

## 1 Introduction

The determination of protein structures stands at the forefront of modern structural biology, providing essential insights into cellular mechanisms, disease processes, and therapeutic development [1]. Understanding the structure of proteins is essential for investigating protein interactions, comprehending pathophysiology, and advancing drug development [2, 3]. Cryo-electron microscopy, or cryo-EM, has emerged as a groundbreaking technology for structure determination, enabling near-atomic resolution imaging of large macromolecular complexes by preserving specimens in their native state through rapid vitrification [4, 5, 6].

A pivotal step in the cryo-EM workflow is protein particle picking, which involves identifying and extracting individual protein particles from micrographs containing thousands of randomly oriented molecules in noisy backgrounds [7]. This process presents substantial technical challenges that directly influence the quality of downstream structural analysis. Cryo-EM micrographs are inherently characterized by extremely low signal-to-noise ratios (SNR), rendering protein particles as subtle contrast variations that are often indistinguishable from background noise, ice contaminations, and various imaging artifacts [8, 9]. The complexity is further compounded by the intrinsic heterogeneity of biological specimens, which may exhibit conformational flexibility, preferred orientations, and structural variability. Additionally, aggregated particles, overlapping structures, and false positives such as ice crystals further complicate reliable particle detection [10, 11, 12]. Therefore, robust and efficient particle picking is essential for ensuring high-resolution cryo-EM structures, as the accuracy and effectiveness of this process have a significant impact on the quality of the resulting 3D density map reconstruction and its resolution.

Traditional particle picking approaches have progressed from fully manual selection to semi-automated template-based methods incorporated in commonly used software packages such as EMAN2 [13], RELION [14], Scipion [15], Dog Picker [16], and APPION [17]. Manual picking, while accurate, is time-consuming and impractical for large datasets [18, 19]. Template-based picking methods involve the generation of 2D reference templates from initial particle subsets, which are then cross-correlated with the entire micrograph through iterative refinement to guide the picking process [1, 18, 19]. Although these approaches have demonstrated effectiveness for well-characterized proteins under favorable signal-to-noise conditions, their performance remains constrained by template quality and representativeness. The inherent limitations of template-based methods include their dependence on iterative user intervention, extensive parameter optimization requirements, and susceptibility to template selection bias. These constraints limit their applicability when applied to new targets or heterogeneous datasets and introduce operator-dependence variability that can compromise reproducibility. Furthermore, templates may inadequately represent variations in particle orientation, size, or imaging conditions, leading to reduced accuracy in complex experimental scenarios [1, 18, 19, 20, 21, 22, 23].

The advancement in deep learning methods has shown great promise in particle picking automation strategies. A number of models, such as APPLE picker [23], DeepPicker [24], AutoCryoPicker [25], Warp [26], CASSPER [27], Topaz [28], CrYOLO [29], and CryoMAE [30] have been developed to improve detection accuracy and reduce manual intervention. Among them, Topaz and CrYOLO, both adopting convolutional neural network (CNN)-based models, remain the most widely used in the cryo-EM community. However, CrYOLO frequently overlooks true protein particles from micrographs, while Topaz demonstrates susceptibility to false positive detection, including ice contaminants and duplicate particles [30, 31, 32]. More recent deep learning approaches, including CryoSegNet [31] and CryoTransformer [32], have shown enhanced performance through sophisticated architectural designs and extensive training procedures. However, these methods typically require large, well-curated training datasets and substantial computational resources for model optimization, making it challenging to apply them in scenarios where limited training data is available [21]. The reliance on extensive training data also raises concerns about generalization to novel particle types or experimental conditions not well-represented in the training datasets. The trade-offs between annotation burden and model adaptability underscore a key gap in Cryo-EM, a paradigm that balances efficient learning with strong flexibility under extreme data scarcity.

This study presents CryoFSL, a data-efficient, few-shot learning framework for protein particle picking in cryo-EM micrographs, leveraging the Segment Anything Model 2 (SAM2) [33], a state-of-the-art vision foundation model. Our approach incorporates lightweight adapter modules into SAM2’s image encoder while maintaining the base model in a frozen state, enabling rapid and effective adaptation to novel protein specimens using as few as five manually annotated micrographs. Although built on a deep learning foundation, CryoFSL’s working mechanism closely resembles template-based methods in practice, offering efficient adaptation to new targets with minimal supervision and no need for large-scale retraining. CryoFSL is specifically designed for the practical settings where current particle-picking methods struggle: **(1) novel or low-resource projects** where only a handful of annotated micrographs are available (e.g., early screening of a new target or small labs without extensive annotation resources); **(2) low signal-to-noise and heterogeneous datasets** where particle contrast varies strongly across micrographs and templates or fully pretrained supervised models fail to generalize; **(3) workflows that prioritize particle quality over raw quantity**, such as downstream projects requiring high-quality reconstruction from fewer and cleaner particles, and **(4) computationally constrained environments** where full model re-training is impractical. By directly addressing these challenges, CryoFSL bridges the gap between accuracy, annotation efficiency, and generalizability, providing a practical solution for both exploratory and large-scale cryo-EM studies.

We tested our framework against traditional template-based methods (EMAN2, RELION, and Scipion) and deep learning approaches (Topaz and CrYOLO), demonstrating superior performance in particle detection across a variety of proteins. While deep learning models like CryoSegNet and CryoTransformer have shown promising results in fully supervised settings, we did not include them in our evaluation due to their dependency on large, curated training datasets and resource-intensive training procedures. These characteristics make them unsuitable for the few-shot, low-annotation regime targeted by CryoFSL. Notably, our method excels in challenging scenarios characterized by low signal-to-noise ratios and significant structural heterogeneity. Our findings highlight the potential of combining foundational models with few-shot learning paradigms to address longstanding challenges in cryo-EM particle picking, offering a scalable, accurate, and annotation-efficient solution for structural biology applications.

## 2 Results

### CryoFSL framework for few-shot protein particle picking

We introduce **CryoFSL**, a parameter-efficient few-shot learning framework for automated protein particle picking in cryo-EM micrographs, utilizing SAM2 enhanced with task-specific adapter modules. As illustrated in Figure 1, input micrographs are first encoded through the frozen SAM2 Hiera-large hierarchical vision transformer encoder to extract multi-scale features, crucial for detecting particles in noisy, low-contrast images. To adapt SAM2 for cryo-EM without full finetuning, lightweight adapter modules are integrated at each encoder stage, comprising a stage-specific unshared linear layer and a shared linear projection that modulate features via residual connections, preserving pretrained knowledge while enabling task-specific adaptation (see section 4). The adapted features are passed to the SAM2 mask decoder to generate a dense segmentation mask identifying potential particle regions. The resulting binary masks undergo comprehensive post-processing through a multi-stage pipeline that combines distance transforms, multi-scale peak detection, and watershed segmentation to extract precise particle coordinates **(Supplementary Algorithm S1)**. The final output includes protein particle coordinates in STAR files, compatible with tools like RELION and CryoSPARC [34] for generating 3D protein density maps, and visually segmented particles marked with circles on the particle regions.

**Figure 1:**
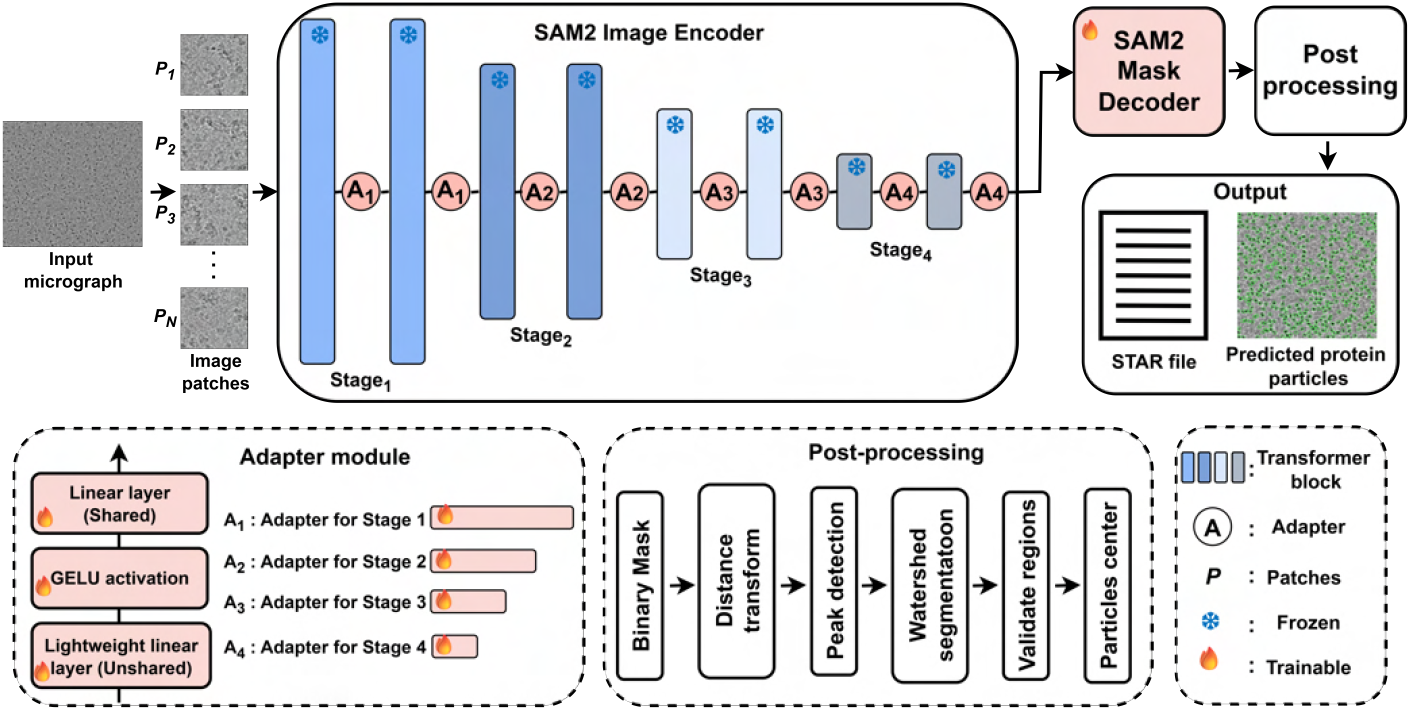
Model architecture of CryoFSL for few-shot cryo-EM protein particle picking. The framework takes a small set of annotated cryo-EM micrographs as input. It utilizes the SAM2 image encoder as its backbone, enhanced with lightweight adapter modules to enable efficient fine-tuning under limited supervision. The encoder is divided into four sequential stages (Stage_1_ to Stage_4_), each containing multiple transformer blocks shown as vertical bars. Adapter modules (*A*_1_ - *A*_4_) inserted between these blocks consist of a shared linear layer, GELU activation, and an unshared lightweight linear layer, enabling targeted adaptation of feature representation while keeping most of the backbone frozen. Frozen and trainable components are visually distinguished to highlight the minimal parameter footprint. The encoded features are passed through the SAM2 mask decoder to generate binary segmentation masks, which undergo a structured post-processing pipeline to generate output. The final outputs are a STAR file (text document) for downstream cryo-EM analysis and a visual map of predicted protein particles. This architecture allows CryoFSL to achieve strong generalization with minimal annotations, leveraging the power of vision foundation models while maintaining adaptability to the heterogeneity and noise in cryo-EM micrographs.

To thoroughly evaluate CryoFSL’s performance, we conducted extensive experiments on the CryoPPP dataset [35], testing 1-shot, 5-shot, and 10-shot scenarios across six diverse proteins with varying morphological complexities from EMPIAR (10028, 10081, 10017, 10093, 10345, and 11056) [36]. We used precision, recall, F1-score, Intersection over Union (IoU), and 3D reconstruction resolution (in Angstroms) to measure particle picking accuracy. To ensure consistency across methods, all baseline models were trained using only five annotated micrographs per protein, matching the few-shot training setup used for CryoFSL, before being tested on the same dataset. Each method was configured according to published best practices, with further details of training and implementation provided in the **Methods** section. The evaluation revealed that CryoFSL uniformly outperformed all baseline models across different scenarios, demonstrating its stability and effectiveness in few-shot settings with minimal labeled data.

### Comparative evaluation of 1-shot, 5-shot, and 10-shot learning for CryoFSL particle picking

We assessed CryoFSL’s few-shot learning capabilities under 1-shot, 5-shot, and 10-shot configurations, where the model was trained using 1, 5, and 10 annotated micrographs per protein, respectively, to investigate the trade-off between annotation effort and model performance. The results are shown in Figure 2. As expected, the 10-shot setup achieved the highest performance, particularly excelling in datasets like 10081, 10093, and 11056. Impressively, the 1-shot scenario revealed CryoFSL’s tolerance for sparse annotations by achieving an acceptable F1-score above 50 % on most proteins despite only using one annotated micrograph.

**Figure 2:**
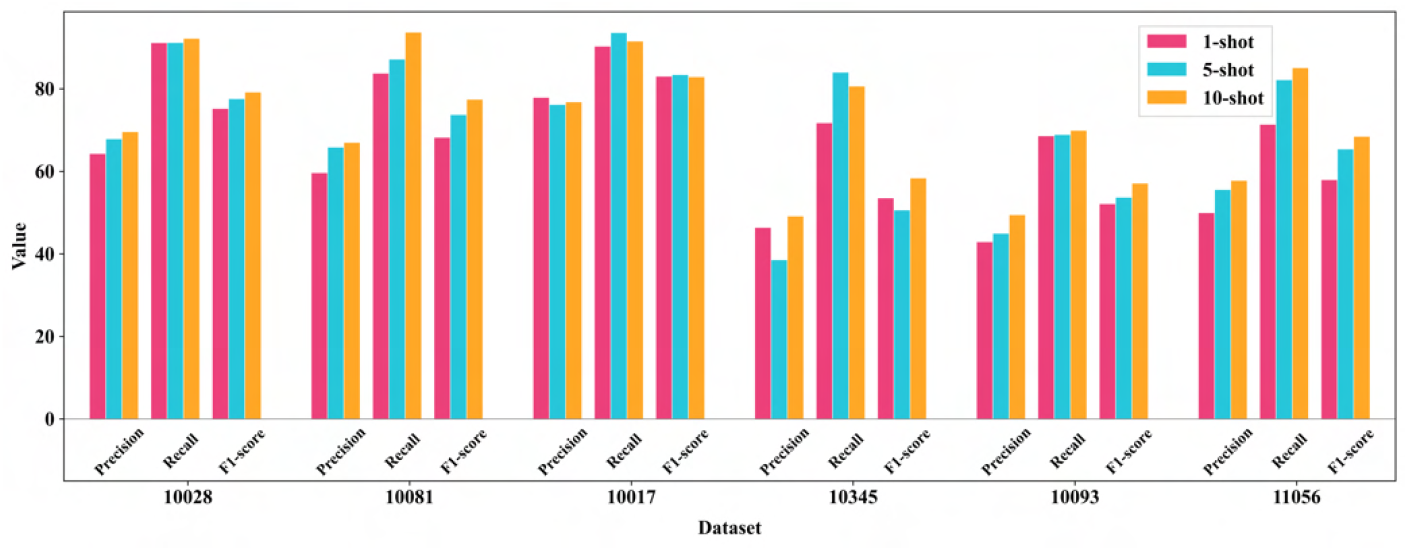
Detailed comparison of CryoFSL’s performance across 1-shot, 5-shot, and 10-shot learning scenarios, evaluated using precision, recall, and F1-score metrics on six diverse cryo-EM protein datasets from the EMPIARs (10028, 10081, 10017, 10345, 10093, and 11056). The x-axis represents those six datasets, and the y-axis displays the numerical value of the performance metrics, ranging from 0 to 100, where higher values indicate better performance. Three few-shot learning scenarios are compared using color-coded bars: 1-shot learning (pink), 5-shot learning (cyan), and 10-shot learning (orange), where each scenario corresponds to the number of annotated micrographs used for training.

The progression from 1-shot to 5-shot yielded greatly enhanced performance, with the 5-shot configuration achieving results remarkably close to 10-shot across all datasets. In certain instances, like EMPIAR-10017 and 10345, the 5-shot CryoFSL even outperformed 10-shot learning with better recall scores. Notably, the marginal gains from 5-shot to 10-shot settings were modest, with 5-shot achieving 90 – 95 % of the maximum attainable performance while requiring only half the annotated data and computational resources. Consequently, we adopted the 5-shot configuration as our optimal training paradigm for all subsequent experiments, as it struck an ideal balance between performance excellence and practical feasibility. This choice supported one of the central goals of our framework: achieving strong generalization with minimal supervision, thereby enabling rapid and scalable deployment in real-world Cryo-EM analysis workflows.

### Systematic comparison of methods using segmentation metrics across diverse cryo-EM datasets in few-shot learning settings

CryoFSL demonstrated excellent overall performance across all six protein targets, achieving the highest average recall (0.845), and F1-score (0.684), as shown in Table 1. In contrast, template-based methods like RELION, EMAN2, and Scipion achieved high recall (e.g., 0.793 for RELION and 0.787 for Scipion) but considerably lower precision (0.352 and 0.367), leading to suboptimal F1-scores of 0.471 and 0.478. Even deep learning methods designed for low-data regimes, such as Topaz and CrYOLO, struggled in challenging cases like EMPIAR 10345, where they achieved F1-scores of just 0.074 and 0.078, respectively. **Supplementary Figure S1** confirms their failure to identify many true particles, reflecting poor adaptability under morphological complexity and low contrast. This suggests that such methods may require more labeled data to achieve reliable results. In contrast, CryoFSL achieved a stable F1-score of 0.528 on the same dataset, leveraging adaptive feature modulation to effectively capture particle characteristics. This resilience with minimal labeled data highlights CryoFSL’s efficiency in few-shot settings, significantly reducing the need for extensive annotations and enhancing reliability for complex cryo-EM micrographs.

**Table 1:** Quantitative evaluation results on six protein datasets from CryoPPP for a few-shot setting. Column 1 lists the EMPIAR IDs, and Column 2 specifies the corresponding protein types. The remaining column reports the evaluation metrics – precision, recall, F1-score and IoU – for each compared method. For each metric, the best performance across methods is highlighted in **bold**. The final row presents the average performance of each method across all six proteins, with the best average also highlighted.

| EMPIAR ID | Protein type | Metrics | CrYOLO | Topaz | EMAN2 | RELION | Scipion | CryoFSL |
| --- | --- | --- | --- | --- | --- | --- | --- | --- |
| 10028 [37] | Ribosome (80S) | Precision | **0.741** | 0.693 | 0.444 | 0.501 | 0.448 | 0.678 |
|  |  | Recall | 0.764 | 0.749 | 0.875 | 0.899 | 0.898 | **0.911** |
|  |  | F1-Score | 0.752 | 0.719 | 0.589 | 0.643 | 0.597 | **0.777** |
|  |  | IoU | 0.602 | 0.562 | 0.417 | 0.474 | 0.426 | **0.636** |
| 10081 [38] | Transport | Precision | 0.641 | 0.590 | 0.353 | 0.377 | 0.365 | **0.658** |
|  |  | Recall | 0.665 | 0.850 | 0.805 | 0.804 | 0.807 | **0.872** |
|  |  | F1-Score | 0.652 | 0.696 | 0.491 | 0.513 | 0.502 | **0.751** |
|  |  | IoU | 0.484 | 0.534 | 0.325 | 0.345 | 0.335 | **0.600** |
| 10017 [39] | β-Galactosidase | Precision | 0.754 | 0.742 | 0.591 | 0.372 | 0.500 | **0.762** |
|  |  | Recall | 0.361 | 0.709 | 0.552 | 0.753 | 0.832 | **0.936** |
|  |  | F1-Score | 0.488 | 0.725 | 0.571 | 0.497 | 0.624 | **0.839** |
|  |  | IoU | 0.322 | 0.567 | 0.399 | 0.331 | 0.454 | **0.724** |
| 10345 [40] | Signaling | Precision | 0.142 | 0.047 | 0.133 | 0.105 | 0.080 | **0.386** |
|  |  | Recall | 0.054 | 0.185 | 0.623 | **0.874** | 0.858 | 0.840 |
|  |  | F1-Score | 0.078 | 0.074 | 0.219 | 0.187 | 0.143 | **0.528** |
|  |  | IoU | 0.041 | 0.039 | 0.123 | 0.103 | 0.078 | **0.359** |
| 10093 [41] | Membrane | Precision | 0.536 | **0.571** | 0.376 | 0.385 | 0.324 | 0.450 |
|  |  | Recall | 0.478 | 0.434 | 0.511 | 0.628 | 0.596 | **0.689** |
|  |  | F1-Score | 0.505 | 0.493 | 0.433 | 0.477 | 0.419 | **0.544** |
|  |  | IoU | 0.338 | 0.327 | 0.276 | 0.314 | 0.266 | **0.373** |
| 11056 [42] | Transport | Precision | **0.670** | 0.624 | 0.507 | 0.372 | 0.482 | 0.556 |
|  |  | Recall | 0.496 | 0.478 | 0.608 | 0.799 | 0.734 | **0.822** |
|  |  | F1-Score | 0.571 | 0.541 | 0.553 | 0.507 | 0.581 | **0.663** |
|  |  | IoU | 0.399 | 0.371 | 0.382 | 0.340 | 0.409 | **0.495** |
| **Average** |  | Precision | 0.581 | 0.545 | 0.401 | 0.352 | 0.367 | **0.582** |
|  |  | Recall | 0.470 | 0.568 | 0.662 | 0.793 | 0.787 | **0.845** |
|  |  | F1-Score | 0.508 | 0.541 | 0.476 | 0.471 | 0.478 | **0.684** |
|  |  | IoU | 0.364 | 0.400 | 0.320 | 0.318 | 0.328 | **0.531** |

Visual comparisons on selected micrographs from EMPIAR-10028, 10081, and 10017 further illustrate these differences (Figure 3). CrYOLO generally under-picks, missing a large number of true particles. Topaz, while capturing most particles from the ground truth (GT), suffers from high redundancy due to frequent overlapping picks. Template-based methods like RELION, Scipion, and EMAN2, on the other hand, exhibit aggressive over-picking behavior, identifying excessive numbers of false positives and including non-particle regions, leading to noisy and less reliable selections. This is especially evident in the magnified regions of Figure 3(b), where multiple overlapping or ambiguous picks are observed. CryoFSL, however, achieves a strong balance: maintaining precise alignment with expert-annotated GT particles while simultaneously detecting additional valid particles overlooked during manual curation. **Supplementary Figures S2-S6** visually reinforce this, highlighting the widespread false detections by competing methods and the clean, uniform particle selections achieved by CryoFSL.

**Figure 3:**
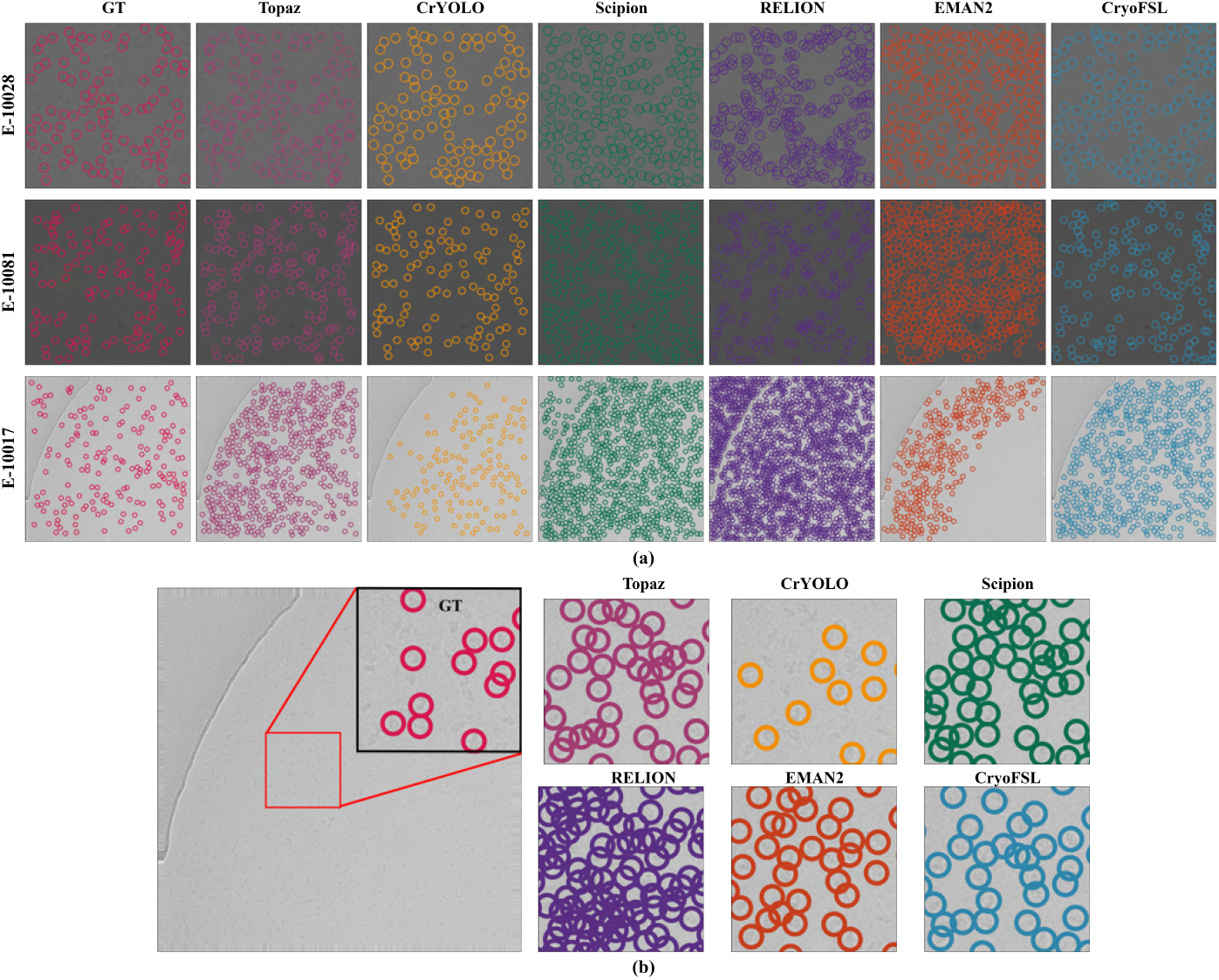
Visual comparison of particle picking across methods and sample micrographs. (a) Sample micrographs from EMPIAR IDs 10028, 10081, and 10017 are shown with particle locations predicted by all competing methods, overlaid alongside the expert-annotated ground truth (GT). Each method’s result is shown as colored circular markers on the micrographs. All models were evaluated on the same set of micrographs to ensure consistency in qualitative comparison.(b) A magnified view of a representative region from a micrograph in EMPIAR 10017 highlights differences in particle localization between the methods. The red box on the full micrograph shows the zoomed region.

### Robustness and failure mode analysis of particle picking methods using segmentation metric distributions

The boxplot analysis in Figure 4 reveals considerable variability in precision and recall distributions across methods, as evidenced by wider interquartile ranges, frequent outliers, and inconsistent medians, particularly for template-based approaches such as RELION and EMAN2. These large spreads and skewed distributions indicate unstable particle picking behavior across different micrographs, often stemming from over-picking in cluttered or noisy regions. The outliers represent instances where template correlation thresholds become either too permissive (generating excessive false positives) or too restrictive (missing true targets), suggesting that fixed-parameter approaches cannot dynamically adapt to the heterogeneous nature of cryo-EM micrographs. This phenomenon is especially pronounced in EMPIARs – 10345 and 10093, where particle diversity and contamination challenge the reproducibility of the templates. The resulting inconsistency is also reflected in Table 1, where high recall is often offset by low precision and F1-score.

**Figure 4:**
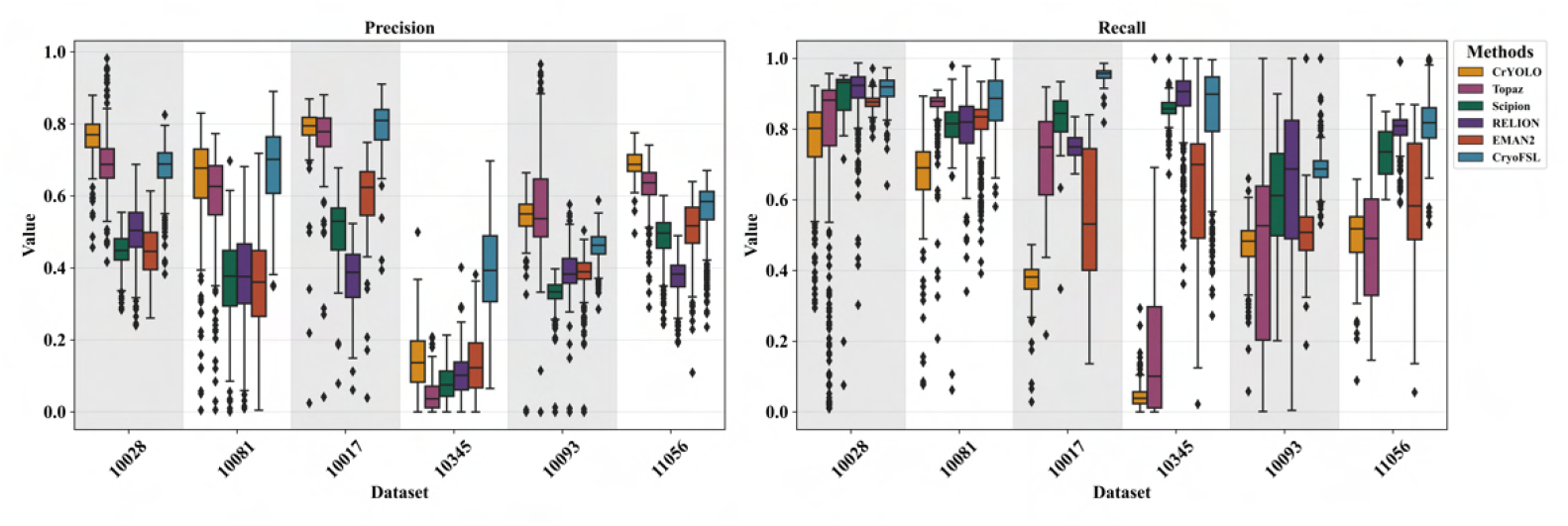
Detailed box plots comparing precision (left) and recall (right) distribution across six methods on six different datasets. The x-axis for both plots represents the six datasets (10028, 10081, 10017, 10345, 10093, and 11056), while the y-axis displays the metric values, ranging from 0 to 1. Each box represents the interquartile range (IQR), the black horizontal line inside each box indicates the median value, and the whiskers extend to the range of non-outlier values. Outliers are plotted as individual points.

In contrast, CryoFSL achieves comparatively tighter distributions across both precision and recall metrics, with fewer outliers and generally higher medians than other methods across most protein datasets. While variability is still observed in some cases, the narrower interquartile ranges highlight more stable behavior that adapts dynamically to varying imaging conditions rather than relying on static decision boundaries. This stability stems from CryoFSL’s adapter modules’ ability to modulate feature representation in response to local micrograph characteristics, effectively learning to distinguish signal from noise patterns that would confound correlation-based approaches. Statistical analysis using Wilcoxon signed-rank tests [43] confirms CryoFSL’s superior performance, with highly significant adjusted p-values (e.g., < 1e^*−*10^ for recall in most cases) and large effect sizes, as detailed in **Supplementary Table S4-S5**. Further, threshold-based analysis shows CryoFSL achieves a 96.6 % recall success rate at the 0.6 threshold and maintains 69.1 % at 0.8, outperforming all baselines in both performance and consistency, as detailed in **Supplementary Figure S7**.

A combined analysis of segmentation behavior across multiple EMPIAR-10345 micrographs **(Supplementary Figure S6 and Supplementary Table S3)** highlights distinct failure modes among methods. CryoFSL maintains particle predictions within a stable range (typically 100-150), reflecting high recall and robustness, with acceptable precision due to detecting valid but unlabeled particles. In contrast, CrYOLO consistently under-picks (as few as 9-26), while Topaz exhibits highly variable behavior, with predicted counts ranging from 5 to 363 depending on micrograph complexity. Template-based methods such as Scipion, RELION, and EMAN2 exhibit aggressive over-picking, achieving high recall but extremely low precision due to genuine false positive detections. Notably, CryoFSL’s lower precision arises from recovery of true but unlabeled particles, fundamentally differing from the false positive-driven precision loss in traditional methods.

### Comparative analysis of particle quantity and 3D density map reconstruction resolution across methods

Table 2 presents a comprehensive evaluation of particle quantity and 3D density map reconstruction quality across methods. Template-based methods routinely selected 2 to 3 times more particles than our approach, yet CryoFSL achieved the best average resolution of 5.33 Å with significantly fewer particles – around 45000 on average. This inverse relationship is particularly striking for challenging proteins like 10345, where CryoFSL’s precise selection (21008 particles) yields exceptional 3.84 Å resolution, outperforming all other methods by greater than 1.44 Å despite their much larger particle sets. High particle numbers are regularly accompanied by elevated resolution value (i.e., lower quality), with RELION reaching 7.36 Å on 10028 despite picking 68055 particles, and EMAN2 yielding 7.26 Å on 10081 with 97234 particles. This observation strongly corroborates our earlier findings from Table 1 and Figure 3, where traditional methods exhibited high recall but low precision and significant performance variability. The superior 3D density recovery of CryoFSL is further demonstrated in Figure 5, which visualizes the density maps for EMPIAR-10345, clearly showing finer structural details and fewer artifacts compared to baseline methods. Additional reconstructions and resolution comparisons for all protein datasets are provided in **Supplementary Figures S8-S9**, affirming CryoFSL’s advantage in delivering cleaner, higher-resolution structures.

**Table 2:** Comparison of 3D reconstruction resolution and particle yield across six cryo-EM protein datasets. For each EMPIAR ID, the number of particles picked (#) and the corresponding 3D resolution (Res., in Å) obtained using each method are reported. The final row shows the average particle count and average resolution for each method. The best resolution (lowest Res. value) for each protein is highlighted in **bold**, indicating superior performance in structural recovery.

| EMPIAR IDs | CrYOLO |  | Topaz |  | EMAN2 |  | RELION |  | Scipion |  | CryoFSL |  |
| --- | --- | --- | --- | --- | --- | --- | --- | --- | --- | --- | --- | --- |
|  | # | Res. | # | Res. | # | Res. | # | Res. | # | Res. | # | Res. |
| 10028 | 26121 | 4.15 | 36112 | **4.08** | 59582 | 4.25 | 68055 | 7.36 | 56591 | 4.13 | 30043 | 4.13 |
| 10081 | 36474 | 7.21 | 56753 | 6.51 | 97234 | 7.26 | 104452 | 8.00 | 85168 | 7.36 | 31799 | **6.16** |
| 10017 | 20793 | 5.98 | 47462 | 5.43 | 48132 | 5.61 | 98263 | 5.58 | 84108 | 5.89 | 45349 | **4.99** |
| 10345 | 4256 | 15.38 | 66247 | 5.28 | 83375 | 5.99 | 118637 | 5.81 | 125693 | 5.77 | 21008 | **3.84** |
| 10093 | 42983 | 6.72 | 55636 | 6.80 | 77593 | 7.34 | 101355 | 7.11 | 86495 | 6.72 | 37513 | **6.47** |
| 11056 | 83713 | 8.09 | 101842 | 7.11 | 183448 | 7.64 | 157005 | 7.40 | 188497 | 6.59 | 106643 | **6.44** |
| **Average** | 35723 | 7.92 | 60675 | 5.86 | 91560 | 6.34 | 107961 | 6.87 | 104425 | 6.07 | 45392 | **5.33** |

**Figure 5:**
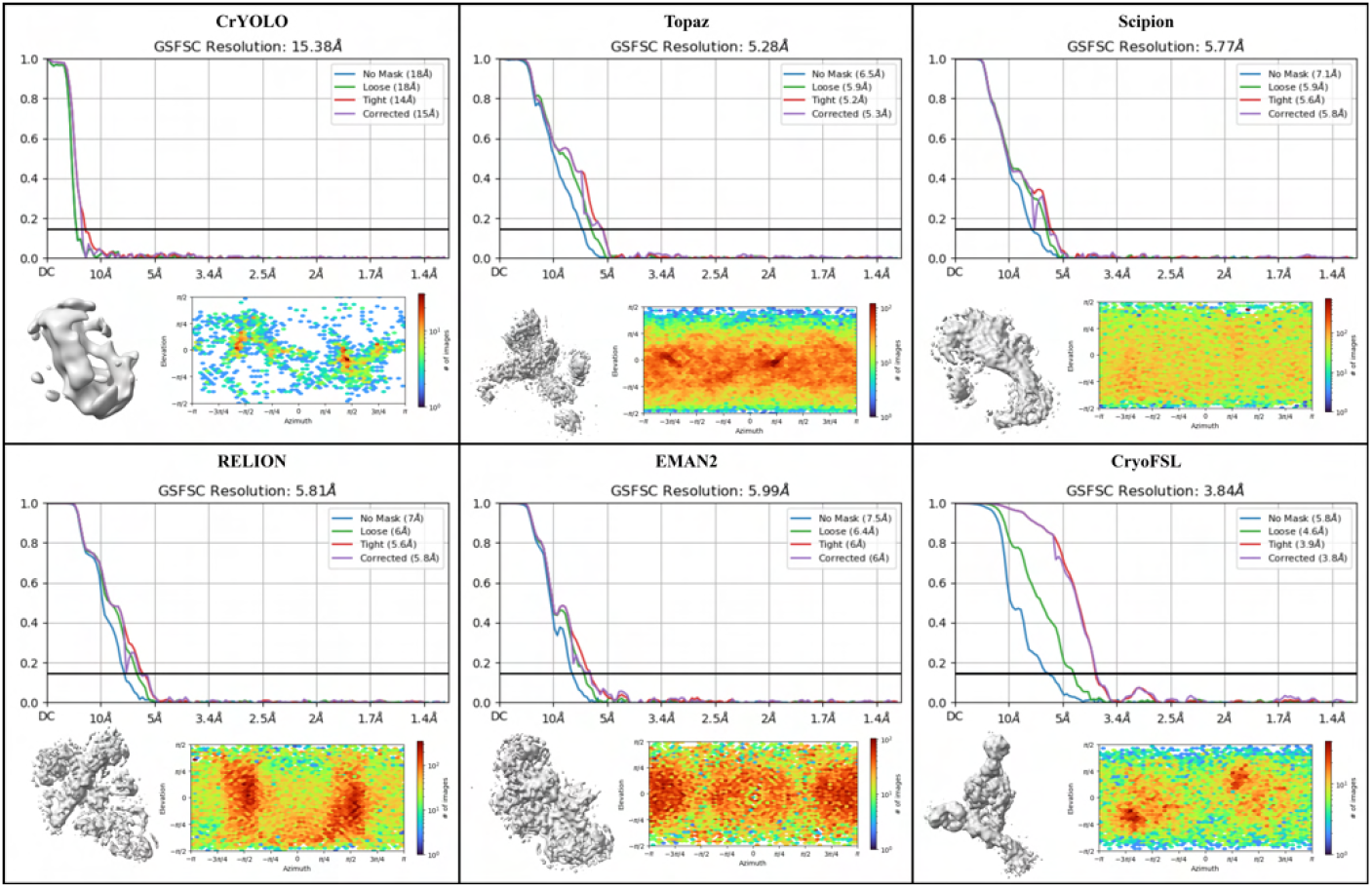
Comparison results for the resolution of the 3D density maps and reconstructed 3D density maps of particles picked by different methods on the EMPIAR-10345 dataset.

To further examine the quality of particles selected by each method, we conducted a progressive reconstruction analysis on EMPIAR 10081 protein, evaluating resolution at 25%, 50%, 75%, and 100 % of the particles picked. Figure 6 reveals that with just 25 % of the particles, CryoFSL achieves a resolution of 7.91 Å, already comparable to or better than the full 100 % sets of EMAN2 (7.26 Å), Scipion (7.36 Å), and RELION (8 Å). As more particles are added, resolution for CryoFSL improves steadily – 6.82 Å at 50%, 6.29 Å at 75%, and 6.16 Å at 100%, but the diminished gain beyond 50 % reflects the robust and high-quality particles across its selection. In contrast, template-based methods like EMAN2 and Scipion exhibit larger performance jumps (e.g., EMAN2 improves from 9.38 Å at 25 % to 7.26 Å at 100%). The higher variability and steeper improvements observed in these methods imply a greater presence of low-quality or noisy particles that dilute reconstruction quality at lower sampling levels.

**Figure 6:**
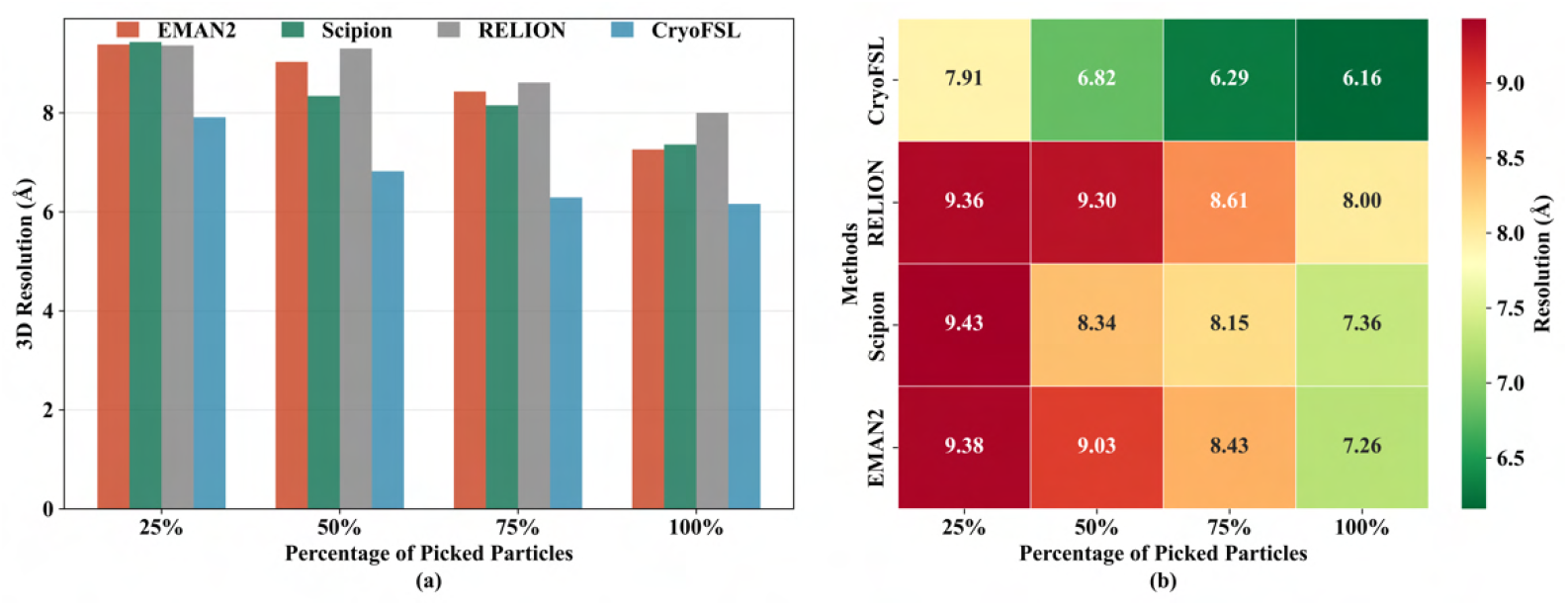
Evaluation of 3D reconstruction resolution across different sampling levels of picked particles for each method using the 10081-protein dataset. (a) Bar plot illustrating the variation in 3D resolution (in Ångström) as a function of the proportion of particles used (25%, 50%, 75%, 100%) for each method. The x-axis represents the percentage of particles used from each method’s picked set, while the y-axis shows the resulting 3D reconstruction resolution in Ångström. The height of each bar reflects the resolution achieved at that sampling level; lower bars indicate better resolution. (b) Heatmap displaying the same data with numerical resolution values annotated in each cell. The x-axis represents the percentage of selected particles used for reconstruction, and the y-axis represents particle picking methods. The color scale encodes resolution values: green indicates superior (lower) resolution, while red denotes poorer (higher) resolution. The color intensity provides an at-a-glance comparison of method performance across sampling levels, with bold annotations for clarity. Together, these visualizations emphasize CryoFSL’s uniformly high-quality particle selection, achieving low resolution values even with a reduced number of particles.

### Evaluating the impact of annotation density on 3D resolution performance across picking methods

To examine performance under limited supervision, we trained each method on five micrographs per protein while varying particle annotations from 10 % to 100%. As shown in Figure 7(a) and **Supplementary Table S2**, CryoFSL demonstrates stable resolution across annotation levels, with only marginal gains at higher supervision. For example, on protein 10017, CryoFSL achieves 5.10 Å at 10 % annotation, nearly matching its 4.99 Å at 100%, and still outperforms EMAN2 (5.61 Å), Scipion (5.89 Å), and RELION (5.58 Å) trained with full annotations. A similar trend holds for protein 10081. This robustness is further quantified in Figure 7(b), where CryoFSL shows the flattest resolution degradation slopes (e.g., −0.00197 Å/ % for 10017), indicating minimal sensitivity to annotation sparsity. In contrast, traditional methods degrade sharply as annotation decreases. These findings highlight CryoFSL’s unique ability to generalize from sparse data.

**Figure 7:**
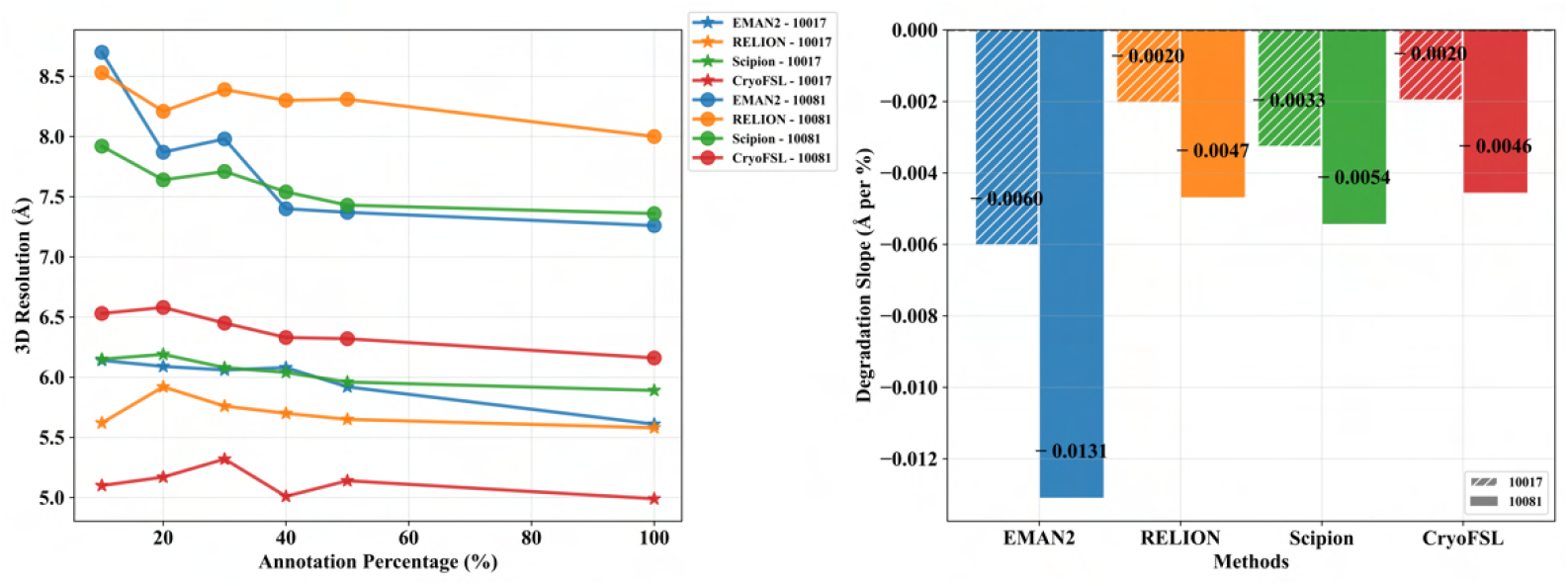
Robustness analysis of cryo-EM protein particle picking methods across varying annotation percentages on two EMPIAR IDs (10017 and 10081).(a) A line plot (left) showing the 3D resolution (Å) achieved by each method with different annotation percentages. The x-axis represents the annotation percentage (%), indicating the proportion of training data available for each method. The y-axis shows the 3D resolution in angstroms (Å), where lower values indicate better performance (i.e., higher resolution structures). Different markers distinguish the two protein datasets: ⋆ (star) for protein 10017 and • (bullet) for protein 10081. Flatter curves indicate greater robustness to reduced training data, while steeper downward slopes suggest higher sensitivity to annotation scarcity. (b) A grouped bar chart (right) displaying the degradation slope (Å per %) for each method–protein combination. The x-axis lists the four approaches, and the y-axis shows the degradation slope in angstroms per percentage point, representing how much the 3D resolution deteriorates for each 1 % reduction in annotation data. Values closer to zero indicate superior robustness (minimal performance degradation with reduced training data). All slopes are negative, which is expected as performance typically degrades with less training data. Bars are grouped by method, with each protein dataset represented by different patterns: diagonal white stripes on a colored background for protein 10017, and solid-colored bars for protein 10081, as indicated in the legend. Numerical values on each bar show the exact slope coefficient. A horizontal dashed line at y = 0 serves as a reference for perfect robustness (no performance degradation).

## 3 Discussion

Cryo-EM particle picking is a crucial step in structural biology, enabling the identification and extraction of individual protein particles from noisy micrographs to reconstruct high-resolution 3D protein structures. This process is inherently challenging due to extremely low signal-to-noise ratios, subtle contrast variations, and the presence of confounding factors like ice contamination and imaging artifacts. Existing approaches, ranging from rigid template matching to fully supervised deep learning models, often fall short in real-world scenarios characterized by protein heterogeneity, limited annotations, and variable imaging conditions. Template-based methods tend to over-pick, misclassifying noise and background as particles due to their reliance on fixed reference shapes, while modern deep learning methods require an extensive labeled dataset and often fail to generalize in low-data or high-complexity regimes. These constraints have created an accessibility barrier that limits the widespread adoption of automated particle picking, particularly for novel protein targets or laboratories with limited computational resources.

To address these challenges, we introduced CryoFSL, a few-shot learning framework that leverages SAM2 enhanced with strategically designed adapter modules to achieve superior particle detection using minimal supervision. Across all evaluations, CryoFSL demonstrated remarkable robustness and flexibility not just by achieving superior average performance metrics, but by maintaining consistency across highly diverse proteins and experimental setups. Unlike conventional methods, CryoFSL was able to identify high-quality particles even when trained with as few as five micrographs, and its performance remained stable across varying levels of annotation density. This level of resilience, especially in heterogeneous or low-contrast micrographs, underscores the strength of its few-shot learning design and its ability to modulate features effectively for novel proteins.

The robustness of CryoFSL stems from its unique architectural design that synergistically combines the foundational visual understanding of SAM2 with task-specific adapter modules integrated across hierarchical encoder stages. This design enables dynamic feature modulation at multiple stages, from local texture patterns distinguishing protein particles from ice crystals to global contextual information resolving ambiguous regions—rather than relying on fixed correlation thresholds that plague traditional methods. While traditional methods exhibit high variance in performance across different micrographs and datasets—evidenced by wide interquartile ranges and numerous outliers—CryoFSL consistently delivers tight performance distributions with minimal variability. Our recall success rate evaluation (96.6 % at 0.6 threshold, 69.1 % at 0.8 threshold) collectively demonstrates unprecedented stability that directly correlates with coherent 3D reconstruction quality. The framework’s exceptional performance under sparse annotation conditions— maintaining near-optimal resolution with only 10 % labeled particles while surpassing deep learning and traditional methods—reveals that the adapter modules efficiently extract maximally informative features rather than memorizing dataset-specific patterns. The visual comparisons further corroborate this: CryoFSL avoids the under-picking of CrYOLO, the redundancy of Topaz, and the aggressive over-picking of template-based methods, achieving a precise balance between recall and precision.

CryoFSL transforms protein identification by implementing what we term *‘intelligent selectivity’*, achieving superior 3D reconstruction resolution (average 5.33 Å) with significantly fewer particles (~45,000) compared to traditional methods that select 2–3 times more particles, yet produce inferior results. This counterintuitive finding reveals that precision in particle selection is far more critical than quantity, exemplified by the EMPIAR-10345 results, where CryoFSL achieved 3.84 Å resolution with only 21,008 particles, while outperforming all baselines by >1.44 Å despite their substantially larger particle counts. The progressive reconstruction analysis further validates this principle, showing that CryoFSL’s particle quality is so high that using only 25 % of selected particles yields resolutions comparable to or better than template-based methods using their complete sets. This transformation occurs because the hierarchical adapter integration enables the framework to distinguish high-quality particles from background noise and artifacts with unprecedented accuracy, effectively mimicking expert microbiologist decision-making through learned feature representation.

The strength of CryoFSL lies in overcoming the fundamental trade-offs that have constrained traditional approaches for decades. Template-based methods achieve high recall through aggressive over-picking but suffer from severely compromised precision due to excessive false positives, while deep learning methods such as Topaz and CrYOLO, when restricted to a few annotated micrographs, exhibited unstable behavior with high variability across proteins. In contrast, CryoFSL maintains consistently high performance across all metrics through its foundational model adaptation strategy. The framework’s few-shot learning capability (optimal performance with just 5 annotated micrographs) addresses the annotation burden that has limited accessibility of automated particle picking, while its parameter-efficient design (lightweight adapters vs. full model retraining) ensures computational feasibility. Most importantly, CryoFSL’s ability to generalize rapidly from minimal examples while maintaining stability across diverse conditions resolves the long-standing tension between automation and accuracy that has plagued the field.

Despite these advances, CryoFSL has limitations that merit consideration. In extremely low-data settings, such as one-shot learning on morphologically complex proteins, the adapter layer may overfit to micrograph-specific noise patterns or sampling artifacts, limiting generalizability. Similarly, very low contrast or severely degraded signal-to-noise ratio (SNR) can render the particle signal indistinguishable from the background. Extreme aggregation, heavy overlaps, or highly irregular particle shapes break assumptions in the post-processing stage (e.g., circularity/area constraints and watershed segmentation), producing merged picks or missed centers. These are fundamentally post-processing failures rather than faults of the adapter strategy itself. Additionally, large domain shifts in imaging conditions— such as unusual microscope settings, uncommon contaminants, or carbon films not represented in the few training micrographs—can leave the frozen backbone’s priors misaligned with task appearance. In such cases, CryoFSL might require modest additional labeled micrographs to adapt effectively. Future enhancements could address these limitations through domain-specific pretraining on larger cryo-EM datasets to improve particle understanding, active learning strategies to guide optimal training example selection, or development of adaptive post-processing algorithms to better handle overlapping particles. Expanding the few-shot framework to downstream tasks like particle classification or heterogeneity analysis could also unlock broader utility in the cryo-EM pipeline.

In conclusion, CryoFSL redefines data efficiency and generalizability in cryo-EM particle picking when few annotated micrographs are available. By combining the strengths of foundational vision models with the flexibility of few-shot adaptation, it bridges the gap between precision and practicality - enabling robust particle identification with minimal annotations, strong generalization across proteins, and higher downstream reconstruction quality. This represents a pivotal step toward scalable and reliable cryo-EM analysis in real-world research environments.

## 4 Methods

### Dataset

We evaluate our approach using the CryoPPP dataset, a large and diverse collection of expertly annotated cryo-EM micrographs curated from the Electron Microscopy Public Image Archive (EMPIAR). CryoPPP encompasses a wide range of protein types, molecular sizes, particle shapes, and imaging conditions, including low signal-to-noise ratios, ice contamination, carbon films, and heterogeneous particle distributions. For our few-shot experiments, we select a representative subset of six proteins from CryoPPP (EMPIAR – 10028, 10081, 10017, 10093, 10345, 11056). These datasets were chosen to reflect variability in particle morphology and micrograph complexity. For each protein, we randomly sample 1, 5, and 10 labeled micrographs to simulate 1-shot, 5-shot, and 10-shot scenarios, respectively. The details of the dataset are presented in Supplementary Table S1.

### Evaluation of particle picking

To quantitatively assess the particle picking performance of CryoFSL and competing approaches, we adopted four widely used metrics: precision, recall, F1-score and Intersection over Union (IoU). Precision measures the proportion of correctly identified particles among all predicted particles, whereas recall measures the proportion of ground truth particles that were correctly detected.

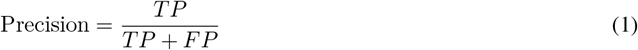

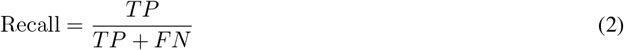

Here, true positive (TP) denotes the predicted particles correctly matched to ground truth, false positive (FP) denotes the predicted particles with no matching ground truth, and false negative (FN) denotes the ground truth particles that were missed. F1-score is the harmonic mean of precision and recall, proving a balanced metric for performance.

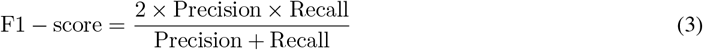

Intersection over Union (IoU) is a measure of the spatial overlap between predicted and ground truth particles.

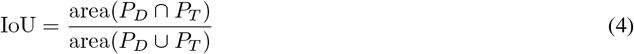

where *P*_*D*_ represents the predicted particles, and *P*_*T*_ represents the ground truth particles.

To further validate the impact of particle picking accuracy on downstream structure determination, we also report the 3D reconstruction resolution obtained using particles picked by each method. This metric reflects the quality of structural recovery and is reported in Angstroms (Å), where lower values indicate higher quality.

### Statistical Analysis

To evaluate the statistical significance of differences between CryoFSL and competing methods, p-values were computed using paired non-parametric tests applied on a per-micrograph basis for each of the six EMPIAR datasets. Specifically, the Wilcoxon signed-rank test was employed to compare paired metric values (precision or recall), assessing the null hypothesis of no difference with CryoFSL as the reference method. Effect sizes were calculated to quantify the magnitude and direction of differences using the rank-biserial correlation [44], computed as 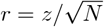, where *z* is the Wilcoxon test statistic and *N* is the number of paired observations.

Given multiple comparisons (5 competitor methods × 6 datasets = 30 tests per metric), raw p-values were adjusted for false discovery using the Benjamini-Hochberg procedure [45] for controlling the false discovery rate (FDR). All statistical computations were implemented in Python using the `scipy.stats.wilcoxon` function for raw p-values and effect sizes, and `statsmodels.stats.multitest.multipletests` with the ‘fdr_bh’ method for adjustments. Final results, including median values, p-values, adjusted p-values, and effect sizes, were tabulated to provide a comprehensive comparison across datasets and methods.

### Overall framework

Overall framework Our framework leverages the Segment Anything Model 2 (SAM2) as the backbone for automated protein particle picking in Cryo-EM micrographs. SAM2 features a Hiera-large hierarchical vision transformer encoder, which efficiently captures multi-scale visual representations through progressive down-sampling and deepening of features. This hierarchical structure enables the model to integrate both local texture and global context - crucial for detecting protein particles in noisy, low-contrast micrographs.

The proposed architecture consists of four main components: the **frozen SAM2 Hiera-large image encoder** for robust feature extraction, novel **lightweight adapter modules** integrated across the encoder’s stages for task-specific adaptation, the **SAM2 mask decoder** for segmentation, and a **post-processing pipeline** to extract particle coordinates from the segmentation mask. Unlike the original SAM [46]/SAM2, CryoFSL omits the prompt encoder and operates without any interactive inputs (e.g., points, boxes, or masks), enabling fully automated particle detection.

### Adapter Integration in SAM2 Image Encoder

To adapt the frozen SAM2 image encoder to the particle picking task, we used parameter-efficient adapter modules following the approach of Chen et al. [47] and He et al. [48], by strategically placing them across all four hierarchical stages of the encoder. Each adapter consists of an unshared linear layer (*L*^unshared^) that captures stage-specific task adaptations, followed by GELU activation for non-linearity, and a shared linear layer (*L*^shared^) that ensures consistent feature dimensionality across stages.

The integration of adapters into the transformer blocks is carefully aligned with the hierarchical structure of the SAM2 image encoder, where each stage processes features of increasing abstraction and semantic complexity. The varying capacities of adapters A_1_ through A_4_ reflect this hierarchical progression: adapter A_1_ operates on the highest spatial resolution features (Stage 1, 144-dimensional embeddings) with two lightweight adapters. At Stage 2, adapter A_2_ processes 288-dimensional embeddings using six adapters. At Stage 3, adapter A_3_ handles 576-dimensional embeddings with 36 adapters—the largest capacity among the stages—consistent with the deeper representation complexity. Finally, adapter A_4_ is integrated into Stage 4, where 1152-dimensional embeddings are projected using four adapters. This architectural design ensures that each hierarchical level benefits from appropriately scaled adaptation capacity, balancing spatial detail in the early stages with semantic richness in the deeper layers.

Let *i* ∈ {1, 2, 3, 4} denote the stage index, and *j* ∈ {0, 1, …, depths[I − 1]} represent the block index within that stage. Let feat_(*i,j*)_ be the input feature to block *j* in stage *i*. Each adapter consists of an unshared linear layer 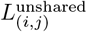 specific to block, GELU activation function σ, and a shared linear layer *L*^shared^ that is common across all adapters. The adapter output for block *j* in stage *i* is formulated as:

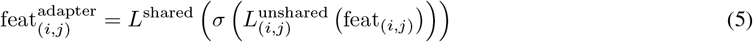

Or, in simplified notation:

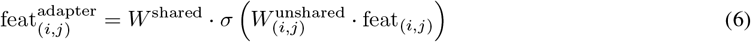

where 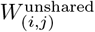 and *W*^shared^ are the learnable weight metrices of the unshared and shared layers, respectively. The task-adapted feature is then added to the original input feature via a residual connection and the final output of block j in stage i is:

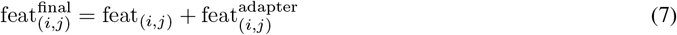

This residual formulation ensures the adapter introduces task-specific modulation without disrupting the pretrained encoder’s underlying representations. Finally, the updated feature 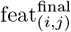 is forwarded through the transformer block BLK_(*i,j*)_ for further processing as:

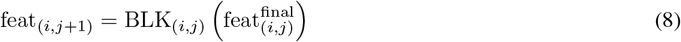

This integration strategy allows each block in the frozen SAM2 encoder to benefit from targeted, low-parameter task adaptation, enabling the model to generalize effectively from a limited number of labeled examples without full fine-tuning of the encoder weights.

### Post-processing and particle localization

To extract accurate particle coordinates from SAM2-generated masks, we employ a robust multi-stage post-processing pipeline combining distance transform, multi-scale peak detection, and watershed segmentation. Geometric filtering based on circularity and area constraints ensures biological plausibility, while a dual-pass strategy enables recovery of closely packed or overlapping particles without duplication. A schematic overview and detailed algorithm are provided in **Supplementary Figure S10**, and **Supplementary Algorithm S1**, respectively.

### Training

CryoFSL was trained using five annotated micrographs from each of the EMPIAR datasets. All micrographs were resized to 1024 × 1024 pixels to align with the input requirements of the SAM2 model. Training was performed using a batch size of 2, the Adam optimizer [49] with a learning rate of 1e^*−*4^, and a maximum of 4000 epochs. The total number of trainable parameters in the model was approximately 3.94 million. For optimization, the balanced binary cross-entropy loss with logits (BCEWithLogitsLoss) was used. During training, the SAM2 image encoder was frozen, and only the adapter module and SAM2 mask decoder were updated. All experiments were implemented in Python 3.11.0, leveraging PyTorch 2.3.0 for model construction and training, and CUDA 11.8 for GPU acceleration.

Template-based methods (EMAN2, RELION, and Scipion) were evaluated using their standard correlation-based workflows. For each protein, templates were generated from the same five labeled micrographs and applied to the test sets with default settings, adjusting only diameter and correlation thresholds when required. No additional fine-tuning or external data were used. Deep learning approaches were likewise trained under identical few-shot configurations. For CrYOLO, we employed PhosaurusNet architecture with an input image of size 768 × 768 and trained for 100 epochs with a learning rate of 1e^*−*4^. Similarly, Topaz used a ResNet backbone, configured with 32 units in the initial layers and batch normalization, trained for 50 epochs with a learning rate of 2e^*−*4^ using the GE-binomial loss. All approaches were then applied to the same test datasets, ensuring a direct and fair comparison with CryoFSL under sparse annotation regimes.

## Author contributions

Study conceptualization (D.X., J.C.); supervision (D.X., J.C.); funding acquisition (D.X., J.C.); methodology and experimental design (B.P.); data preparation and formal analysis (B.P., R.G., A.D.); initial manuscript (B.P.); validation and visualization (B.P., R.G., A.D.). All authors contributed to the revision of the manuscript and have approved the final version.

## Data and Code Availability

The dataset for this study is available on https://github.com/BioinfoMachineLearning/cryoppp. The source code is available at https://github.com/biplabpoudel25/CryoFSL.

