## Supplementary Material for "CryoFSL: An Annotation-Efficient, Few-Shot Learning Framework for Robust Protein Particle Picking in Cryo-EM Micrographs"

**Supplementary Table S1:** An overview of EMPIAR IDs from CryoPPP used for evaluation of CryoFSL (* represents theoretical weight of the proteins)

| SN | EMPIAR ID | Type of Protein | Image Size | Total Structure Weight (kDa) | Number of Images |
|---|---|---|---|---|---|
| 1 | 10028 | Ribosome (80S) | (4096, 4096) | 2135.89 | 300 |
| 2 | 10081 | Transport Protein | (3710, 3838) | 298.57 | 300 |
| 3 | 10345 | Signaling Protein | (3838, 3710) | 244.68 | 295 |
| 4 | 11056 | Transport Protein | (5760, 4092) | 88.94 | 305 |
| 5 | 10093 | Membrane Protein | (3838, 3710) | 779.4 | 295 |
| 6 | 10017 | $\beta$-galactosidase | (4096, 4096) | 450* | 84 |
| | | | | **Total** | **1,879** |

**Supplementary Table S2:** Efficiency of manually annotated particles on 3D resolution for sample proteins using template-based methods and CryoFSL.

| EMPIAR ID | % of Manual Annotation | Number of Particles | EMAN2 | RELION | Scipion | CryoFSL |
|---|---|---|---|---|---|---|
| 10017 | 10% | 170 | 6.14 | 5.62 | 6.15 | 5.10 |
| | 20% | 341 | 6.09 | 5.92 | 6.19 | 5.17 |
| | 30% | 513 | 6.06 | 5.76 | 6.08 | 5.32 |
| | 40% | 684 | 6.08 | 5.70 | 6.04 | 5.01 |
| | 50% | 857 | 5.92 | 5.65 | 5.96 | 5.14 |
| | 100% | 1715 | 5.61 | 5.58 | 5.89 | 4.99 |
| 10081 | 10% | 59 | 8.70 | 8.53 | 7.92 | 6.53 |
| | 20% | 120 | 7.87 | 8.21 | 7.64 | 6.58 |
| | 30% | 180 | 7.98 | 8.39 | 7.71 | 6.45 |
| | 40% | 241 | 7.40 | 8.30 | 7.54 | 6.33 |
| | 50% | 303 | 7.37 | 8.31 | 7.43 | 6.32 |
| | 100% | 607 | 7.26 | 8.00 | 7.36 | 6.16 |

For each protein sample, we investigated the effect of varying the proportion of manually annotated particles on the achievable 3D resolution. The "% of manual annotation" column does not represent the fraction of total training data used; rather, it indicates the fraction of annotated particles selected from each micrograph within the CryoPPP dataset. Specifically, for every micrograph, we subsampled 10%, 20%, 30%, 40%, 50%, and a complete set of high-confidence annotated particles, and then combined these across five micrographs per protein to construct the training or template generation sets. For example, in the case of EMPIAR-10017, selecting 10% of particles corresponds to a total of 170 particles aggregated from the five micrographs (not 170 particles per micrograph). This subsampling strategy allowed us to examine how the resolution scales with increasingly larger sets of particles while keeping the number of micrographs fixed. The resulting particle counts are reported in the "Number of particles" column, which served either as the input for training CryoFSL or as the reference templates for conventional template-based methods. Doing so ensured a fair comparison across methods under varying annotation efficiencies.

**Supplementary Table S3:** Quantitative performance of particle picking across representative micrographs from EMPIAR-10345. Each row reports the expert-curated ground truth (GT) particle count, alongside predicted particle numbers (Pred.), precision (Prec), and recall (Recall) for different methods. The table illustrates the variability in prediction behaviors at the micrograph level, complementing the global statistical analyses presented in the main manuscript.

| Micrograph name | | CrYOLO | | | Topaz | | | Scipion | | | RELION | | | EMAN2 | | | CryoFSL | | |
|---|---|---|---|---|---|---|---|---|---|---|---|---|---|---|---|---|---|---|---|
| | GT | Pred. | Prec | Recall | Pred. | Prec | Recall | Pred. | Prec | Recall | Pred. | Prec | Recall | Pred. | Prec | Recall | Pred. | Prec | Recall |
| 18jam15a_0176_ali_DW | 39 | 26 | 0.2017 | 0.1327 | 83 | 0.01437 | 0.02531 | 653 | 0.06039 | 0.8694 | 709 | 0.0654 | 0.9153 | 104 | 0.12346 | 0.314797 | 134 | 0.2634 | 0.9728 |
| 18jam15a_0185_ali_DW | 16 | 14 | 0.0492 | 0.0408 | 363 | 0.00613 | 0.09916 | 646 | 0.02612 | 0.8959 | 185 | 0.07648 | 0.79825 | 147 | 0.027868 | 0.238854 | 95 | 0.1682 | 0.8361 |
| 18jam15a_0205_ali_DW | 29 | 12 | 0.11761 | 0.04715 | 23 | 0.00961 | 0.00686 | 683 | 0.04242 | 0.84321 | 756 | 0.046 | 0.895 | 156 | 0.124418 | 0.640653 | 112 | 0.2459 | 0.9551 |
| 18jam15a_0225_ali_DW | 29 | 15 | 0.10899 | 0.05653 | 52 | 0.05997 | 0.10276 | 669 | 0.04652 | 0.9141 | 958 | 0.04258 | 0.9533 | 432 | 0.061367 | 0.82755 | 118 | 0.2401 | 0.9949 |
| 18jam15a_0230_ali_DW | 19 | 13 | 0.00013 | 0.01 | 7 | 0.04799 | 0.01768 | 681 | 0.02967 | 0.8943 | 567 | 0.03664 | 0.89792 | 335 | 0.041677 | 0.661564 | 112 | 0.1436 | 0.9744 |
| 18jam15a_0231_ali_DW | 24 | 14 | 0.05117 | 0.03006 | 5 | 0.01957 | 0.00411 | 641 | 0.0358 | 0.8329 | 532 | 0.05091 | 0.92507 | 421 | 0.046974 | 0.757085 | 126 | 0.179 | 0.9335 |
| 18jam15a_0249_ali_DW | 17 | 17 | 0.0819 | 0.0819 | 133 | 0.02642 | 0.1841 | 692 | 0.0248 | 0.8422 | 281 | 0.046 | 0.6887 | 274 | 0.015231 | 0.222002 | 95 | 0.1631 | 0.9791 |
| 18jam15a_0264_ali_DW | 22 | 15 | 0.03221 | 0.02196 | 199 | 0.00331 | 0.02218 | 658 | 0.0331 | 0.83659 | 650 | 0.0416 | 0.9668 | 223 | 0.05107 | 0.483323 | 106 | 0.19309 | 0.9462 |
| 18jam15a_0281_ali_DW | 23 | 16 | 0.04904 | 0.03335 | 15 | 0.0013 | 0.0021 | 674 | 0.0344 | 0.8545 | 753 | 0.03706 | 0.9046 | 267 | 0.064406 | 0.689721 | 123 | 0.1665 | 0.989 |
| 18jam15a_0282_ali_DW | 20 | 9 | 0.11127 | 0.05032 | 11 | 0.031 | 0.1634 | 663 | 0.02885 | 0.82097 | 622 | 0.03537 | 0.8886 | 436 | 0.031198 | 0.624682 | 97 | 0.1865 | 0.9961 |
| 18jam15a_0293_ali_DW | 33 | 20 | 0.12288 | 0.0751 | 14 | 0.0051 | 0.1305 | 667 | 0.0494 | 0.8598 | 835 | 0.0525 | 0.9666 | 447 | 0.057713 | 0.717525 | 162 | 0.1784 | 0.9571 |
| 18jam15a_0295_ali_DW | 39 | 9 | 0.13512 | 0.03074 | 18 | 0.0196 | 0.0054 | 660 | 0.0617 | 0.8853 | 772 | 0.06466 | 0.9504 | 142 | 0.1226 | 0.416327 | 146 | 0.1959 | 0.9793 |
| 18jam15a_0309_ali_DW | 36 | 22 | 0.16515 | 0.10139 | 24 | 0.01437 | 0.0091 | 645 | 0.0563 | 0.868 | 698 | 0.0622 | 0.9245 | 278 | 0.108209 | 0.789765 | 146 | 0.18821 | 0.9814 |
| 18jam15a_0313_ali_DW | 12 | 11 | 0.08121 | 0.07502 | 283 | 0.01362 | 0.2383 | 680 | 0.017 | 0.828 | 342 | 0.0344 | 0.8824 | 281 | 0.019406 | 0.412907 | 46 | 0.19182 | 0.97442 |

To further illustrate the robustness of particle picking methods, we examined 14 representative micrographs from EMPIAR-10345, reporting ground truth (GT) particle count, predicted counts, precision, and recall. Unlike aggregate evaluations that average over many micrographs, this micrograph-level analysis further highlights method-specific behaviors under varying particle densities and image conditions. An important observation from this table is that methods differ not only in aggregate performance but also in consistency across micrographs. For instance, CryoFSL exhibits a narrow-predicted particle range, remaining stable even when ground truth particle counts vary. In contrast, methods like Topaz, EMAN2, and RELION show large fluctuations in predicted counts relative to GT, indicating sensitivity to micrograph-specific factors such as noise patterns or particle clustering. For template-based methods, this manifests as uniformly inflated counts across nearly all micrographs, while Topaz alternates between extreme under-picking (as few as 5, 7) and over-picking (200, 300), reflecting instability in thresholding or learned representation. CrYOLO tends to scale its detections proportionally with particle density but systematically under-recovers absolute particle numbers in low SNR regions, which explains its low recall despite reasonable proportionality with GT. These divergent failure modes have practical consequences: aggressive over-picking from template methods introduces false positives that can degrade downstream alignment and classification, while under-picking methods risk losing true particle diversity needed for high-resolution reconstructions.

**Supplementary Table S4:** Precision comparison of particle picking methods across EMPIAR datasets using Wilcoxon tests. The table reports median precision from CryoFSL (focal) and competing methods, along with $p$-values, effect sizes, and FDR-adjusted $p$-values.

| Dataset | Competitor | Test | $n$ | Median_Focal | Median_Comp | $p$-value | Effect Size | Adj. $p$-value |
|---|---|---|---|---|---|---|---|---|
| 10017 | CrYOLO | Wilcoxon | 79 | 0.80486 | 0.79467 | 0.55103 | 0.18987 | 0.55103 |
| | Topaz | Wilcoxon | 79 | 0.80486 | 0.78428 | 0.044065 | 0.24051 | 0.047212 |
| | Scipion | Wilcoxon | 79 | 0.80486 | 0.52516 | 1.502e-14 | 0.94937 | 1.7331e-14 |
| | RELION | Wilcoxon | 79 | 0.80486 | 0.37835 | 1.289e-14 | 0.94937 | 1.6113e-14 |
| | EMAN2 | Wilcoxon | 79 | 0.80486 | 0.62842 | 1.4457e-14 | 0.92405 | 1.7331e-14 |
| 10028 | CrYOLO | Wilcoxon | 293 | 0.68852 | 0.77012 | 8.4968e-50 | -1.00000 | 3.1863e-49 |
| | Topaz | Wilcoxon | 293 | 0.68852 | 0.68914 | 0.054719 | 0.00341 | 0.056606 |
| | Scipion | Wilcoxon | 293 | 0.68852 | 0.45003 | 8.4968e-50 | 1.00000 | 3.1863e-49 |
| | RELION | Wilcoxon | 293 | 0.68852 | 0.50424 | 8.4968e-50 | 1.00000 | 3.1863e-49 |
| | EMAN2 | Wilcoxon | 293 | 0.68852 | 0.44749 | 8.4968e-50 | 1.00000 | 3.1863e-49 |
| 10081 | CrYOLO | Wilcoxon | 295 | 0.68893 | 0.67734 | 3.3951e-08 | 0.32881 | 3.7724e-08 |
| | Topaz | Wilcoxon | 295 | 0.68893 | 0.62652 | 9.35e-39 | 0.70847 | 1.3357e-38 |
| | Scipion | Wilcoxon | 295 | 0.68893 | 0.37692 | 4.0001e-50 | 1.00000 | 3.1863e-49 |
| | RELION | Wilcoxon | 295 | 0.68893 | 0.37310 | 4.0825e-50 | 0.99322 | 3.1863e-49 |
| | EMAN2 | Wilcoxon | 295 | 0.68893 | 0.35851 | 4.2961e-50 | 0.98644 | 3.1863e-49 |
| 10093 | CrYOLO | Wilcoxon | 285 | 0.46324 | 0.54995 | 1.8433e-48 | -0.97895 | 3.6866e-48 |
| | Topaz | Wilcoxon | 285 | 0.46324 | 0.53729 | 5.0293e-41 | -0.75439 | 7.9409e-41 |
| | Scipion | Wilcoxon | 285 | 0.46324 | 0.33493 | 1.7302e-48 | 1.00000 | 3.6866e-48 |
| | RELION | Wilcoxon | 285 | 0.46324 | 0.38330 | 4.8601e-39 | 0.81053 | 7.2902e-39 |
| | EMAN2 | Wilcoxon | 285 | 0.46324 | 0.39086 | 5.6256e-48 | 0.97193 | 1.0548e-47 |
| 10345 | CrYOLO | Wilcoxon | 291 | 0.39076 | 0.13676 | 2.2891e-49 | 0.97938 | 5.2825e-49 |
| | Topaz | Wilcoxon | 291 | 0.39076 | 0.03683 | 1.8049e-49 | 1.00000 | 4.5122e-49 |
| | Scipion | Wilcoxon | 291 | 0.39076 | 0.07499 | 1.8049e-49 | 1.00000 | 4.5122e-49 |
| | RELION | Wilcoxon | 291 | 0.39076 | 0.10302 | 1.8049e-49 | 1.00000 | 4.5122e-49 |
| | EMAN2 | Wilcoxon | 291 | 0.39076 | 0.12292 | 1.8049e-49 | 1.00000 | 4.5122e-49 |
| 11056 | CrYOLO | Wilcoxon | 299 | 0.58650 | 0.68749 | 8.8662e-51 | -1.00000 | 2.6599e-49 |
| | Topaz | Wilcoxon | 299 | 0.58650 | 0.63838 | 1.2757e-45 | -0.83946 | 2.1261e-45 |
| | Scipion | Wilcoxon | 299 | 0.58650 | 0.49802 | 1.1612e-17 | 0.74582 | 1.5834e-17 |
| | RELION | Wilcoxon | 299 | 0.58650 | 0.38330 | 1.3175e-47 | 0.75251 | 2.3250e-47 |
| | EMAN2 | Wilcoxon | 299 | 0.58650 | 0.51832 | 1.2994e-15 | 0.58528 | 1.6948e-15 |

**Supplementary Table S5:** Recall comparison of particle picking methods across EMPIAR datasets using Wilcoxon tests. The table reports median precision from CryoFSL (focal) and competing methods, along with $p$-values, effect sizes, and FDR-adjusted $p$-values.

| Dataset | Competitor | Test | $n$ | Median_Focal | Median_Comp | $p$-value | Effect Size | Adj. $p$-value |
|---|---|---|---|---|---|---|---|---|
| 10017 | CrYOLO | Wilcoxon | 79 | 0.9557 | 0.38146 | 1.1491E-14 | 1 | 1.8852E-14 |
| | Topaz | Wilcoxon | 79 | 0.9557 | 0.71639 | 1.3915E-14 | 0.94937 | 2.0873E-14 |
| | Scipion | Wilcoxon | 79 | 0.9557 | 0.84421 | 3.1929E-12 | 0.92405 | 4.5613E-12 |
| | RELION | Wilcoxon | 79 | 0.9557 | 0.74916 | 3.4224E-12 | 0.92405 | 4.6669E-12 |
| | EMAN2 | Wilcoxon | 79 | 0.9557 | 0.48612 | 1.1940E-14 | 0.97468 | 1.8852E-14 |
| 10028 | CrYOLO | Wilcoxon | 293 | 0.9187 | 0.80612 | 8.4968E-50 | 1 | 8.4968E-49 |
| | Topaz | Wilcoxon | 293 | 0.9187 | 0.8817 | 8.3895E-39 | 0.73379 | 2.0974E-38 |
| | Scipion | Wilcoxon | 293 | 0.9187 | 0.93211 | 3.5825E-01 | -0.06485 | 3.7061E-01 |
| | RELION | Wilcoxon | 293 | 0.9187 | 0.92361 | 6.4858E-01 | -0.04437 | 6.4858E-01 |
| | EMAN2 | Wilcoxon | 293 | 0.9187 | 0.87667 | 5.4823E-36 | 0.75427 | 1.2652E-35 |
| 10081 | CrYOLO | Wilcoxon | 295 | 0.8848 | 0.69059 | 4.3845E-50 | 0.98644 | 6.5768E-49 |
| | Topaz | Wilcoxon | 295 | 0.8848 | 0.87729 | 1.9602E-01 | 0.11186 | 2.1002E-01 |
| | Scipion | Wilcoxon | 295 | 0.8848 | 0.81501 | 5.5174E-33 | 0.66102 | 1.1823E-32 |
| | RELION | Wilcoxon | 295 | 0.8848 | 0.81982 | 4.9136E-18 | 0.45085 | 9.2131E-18 |
| | EMAN2 | Wilcoxon | 295 | 0.8848 | 0.8339 | 5.6711E-18 | 0.45763 | 1.0008E-17 |
| 10093 | CrYOLO | Wilcoxon | 285 | 0.6875 | 0.48477 | 5.3573E-48 | 0.98947 | 2.2960E-47 |
| | Topaz | Wilcoxon | 285 | 0.6875 | 0.52658 | 8.7489E-41 | 0.75789 | 2.3861E-40 |
| | Scipion | Wilcoxon | 285 | 0.6875 | 0.61673 | 5.9073E-12 | 0.25614 | 7.7052E-12 |
| | RELION | Wilcoxon | 285 | 0.6875 | 0.68713 | 1.5259E-02 | 0.02807 | 1.7607E-02 |
| | EMAN2 | Wilcoxon | 285 | 0.6875 | 0.50705 | 5.3573E-48 | 0.98947 | 2.2960E-47 |
| 10345 | CrYOLO | Wilcoxon | 291 | 0.9027 | 0.03868 | 1.0302E-48 | 0.98282 | 6.1811E-48 |
| | Topaz | Wilcoxon | 291 | 0.9027 | 0.09484 | 1.4939E-47 | 0.8866 | 5.6021E-47 |
| | Scipion | Wilcoxon | 291 | 0.9027 | 0.85865 | 6.0934E-02 | 0.32302 | 6.7704E-02 |
| | RELION | Wilcoxon | 291 | 0.9027 | 0.90623 | 4.0027E-03 | -0.09622 | 5.0033E-03 |
| | EMAN2 | Wilcoxon | 291 | 0.9027 | 0.70119 | 6.7193E-42 | 0.83849 | 2.0158E-41 |
| 11056 | CrYOLO | Wilcoxon | 299 | 0.8180 | 0.51864 | 8.8662E-51 | 1 | 2.6599E-49 |
| | Topaz | Wilcoxon | 299 | 0.8180 | 0.48439 | 1.5624E-49 | 0.9398 | 1.1718E-48 |
| | Scipion | Wilcoxon | 299 | 0.8180 | 0.73528 | 8.3986E-32 | 0.58528 | 1.6797E-31 |
| | RELION | Wilcoxon | 299 | 0.8180 | 0.81038 | 4.2033E-03 | 0.10368 | 5.0439E-03 |
| | EMAN2 | Wilcoxon | 299 | 0.8180 | 0.57915 | 6.7055E-45 | 0.8194 | 2.2352E-44 |

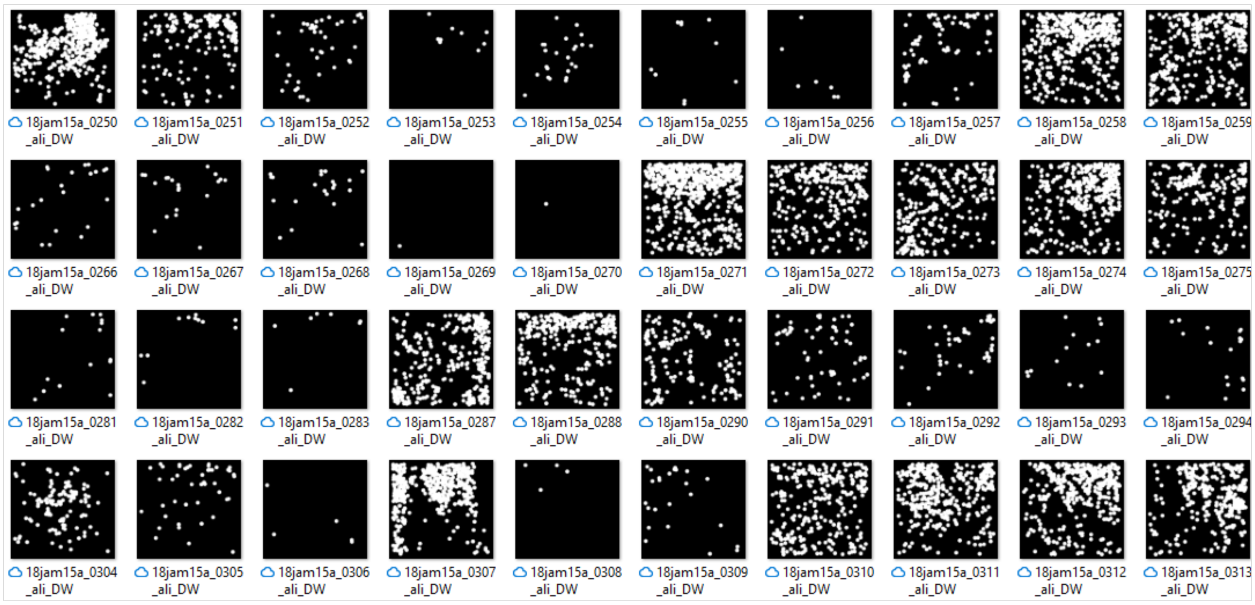


**(a)**

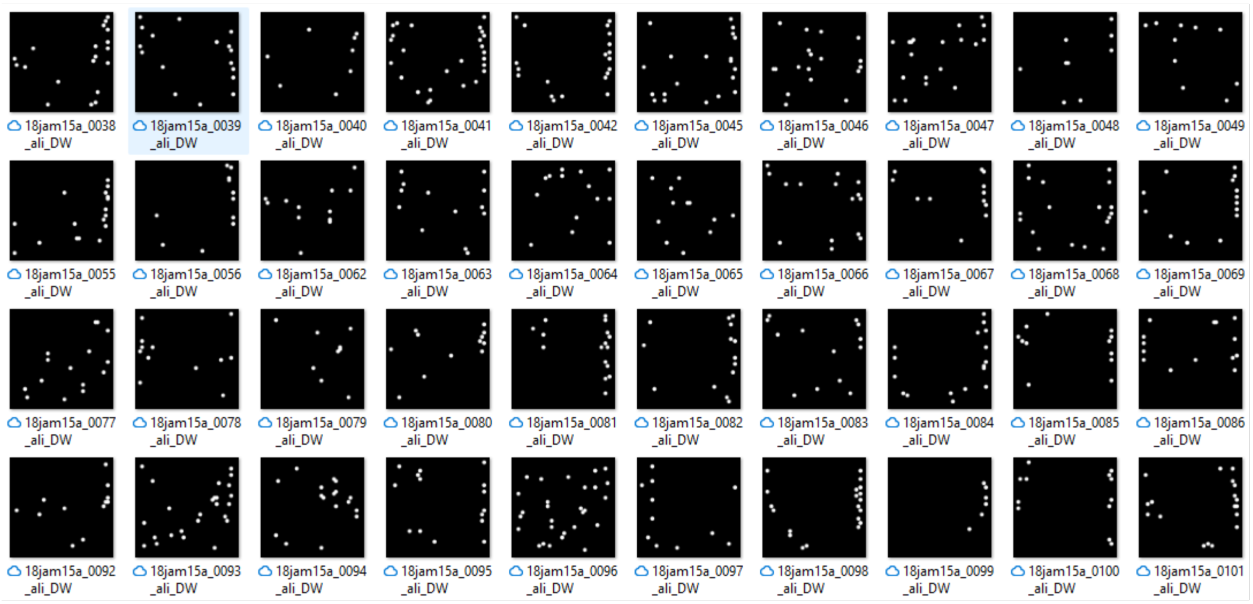


**(b)**

**Supplementary Figure S1:** Visualization of binary mask predictions for protein particles on EMPIAR-10345 micrographs, comparing (a) Topaz and (b) CrYOLO methods. (a) displays Topaz's output, where binary masks (white regions) highlight detected particles, revealing inconsistent performance: over-picking is evident in some micrographs with excessive mask overlaps, while under-picking occurs in others, missing true particles due to low sensitivity in noisy conditions. (b) presents CrYOLO's output, with a binary mask consistently showing under-picking across nearly all sample micrographs, underrepresenting the true number of protein particles, and leaving significant areas undetected. Both figures illustrate the limitations of these methods on the challenging EMPIAR-10345 dataset, characterized by low signal-to-noise ratios and structural heterogeneity.

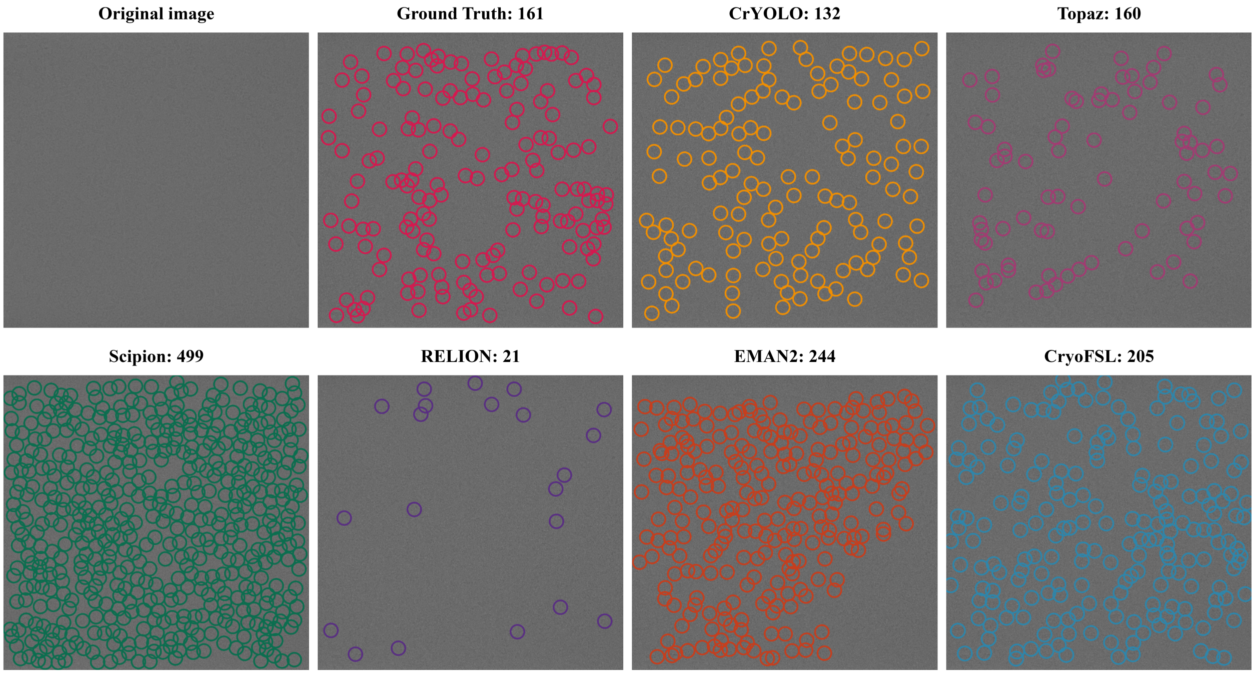


**Supplementary Figure S2:** Visualization of particle picking on EMPIAR-10093 using different methods with their particle counts.

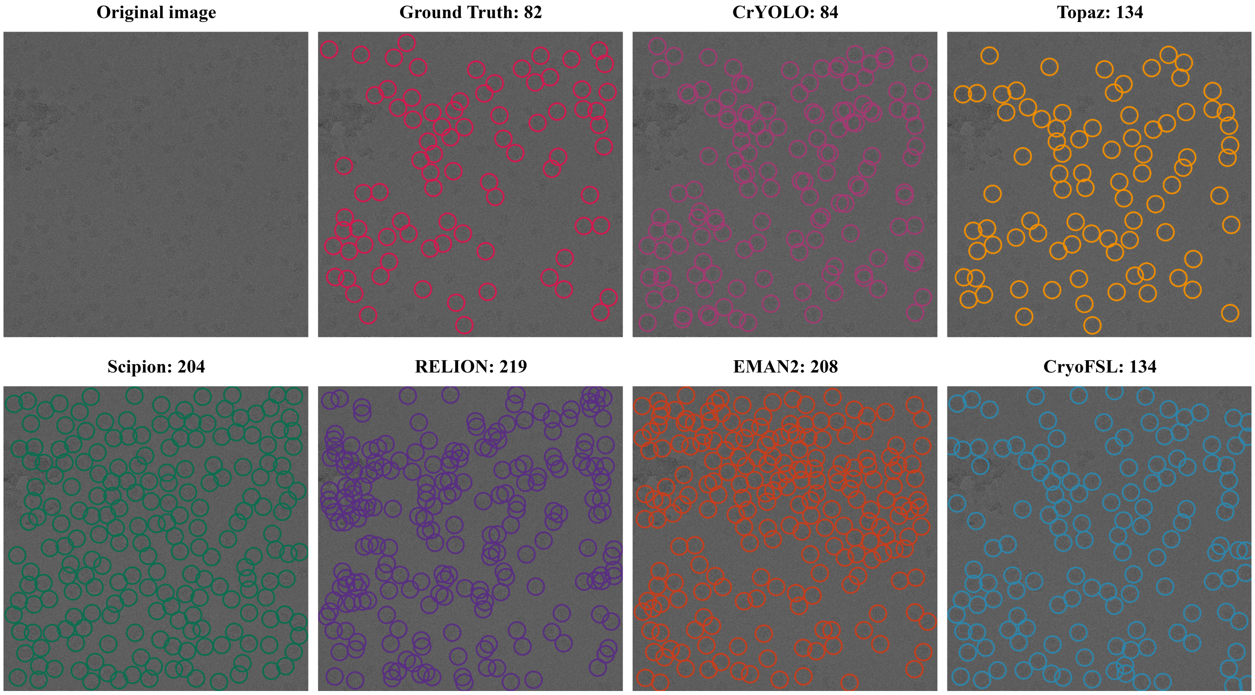


**Supplementary Figure S3:** Visualization of particle picking on EMPIAR-10028 using different methods with their particle counts.

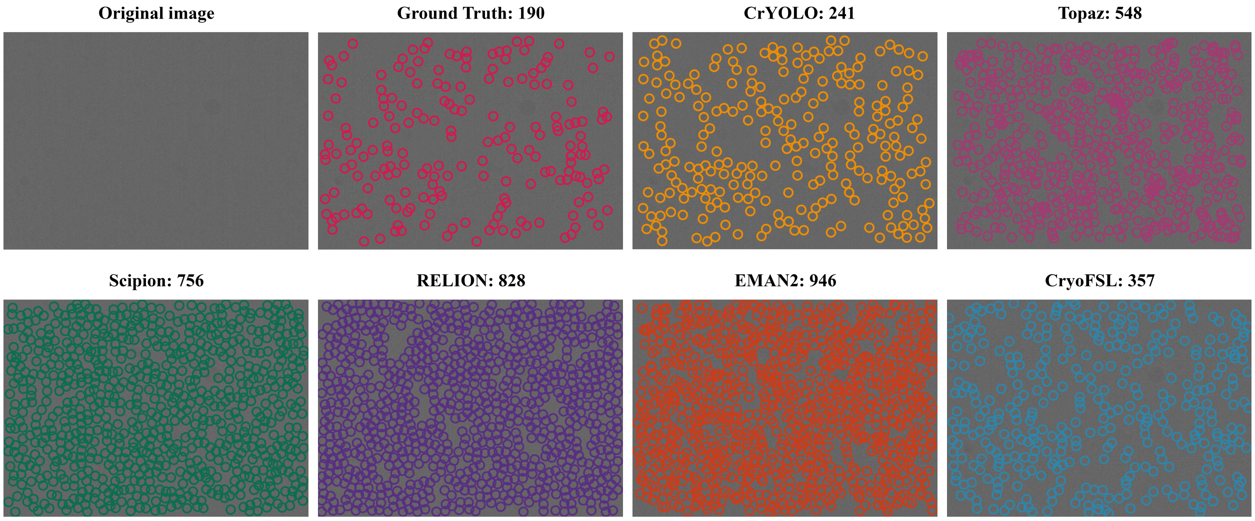


**Supplementary Figure S4:** Visualization of particle picking on EMPIAR-11056 using different methods with their particle counts.

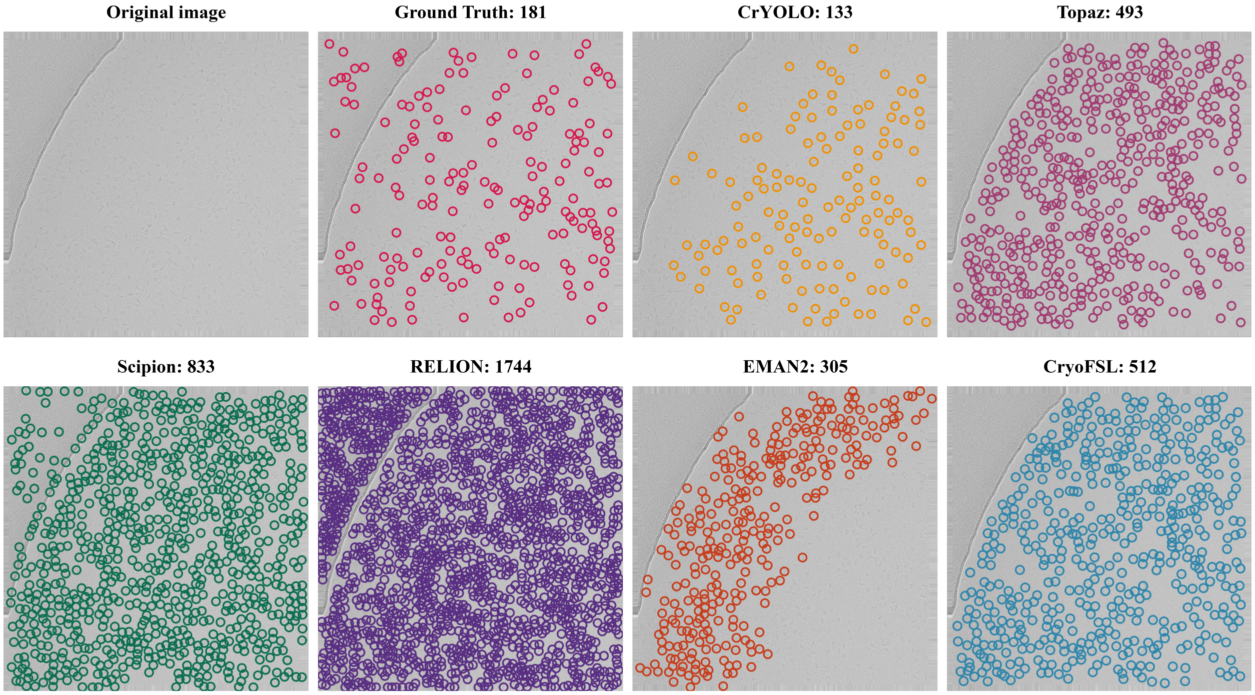


**Supplementary Figure S5:** Visualization of particle picking on EMPIAR-10017 using different methods with their particle counts.

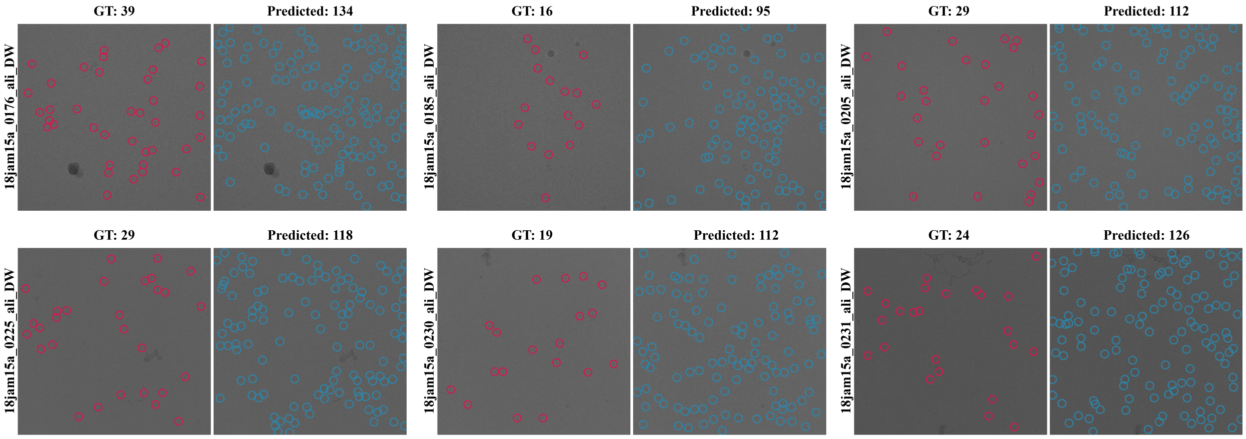


(a)

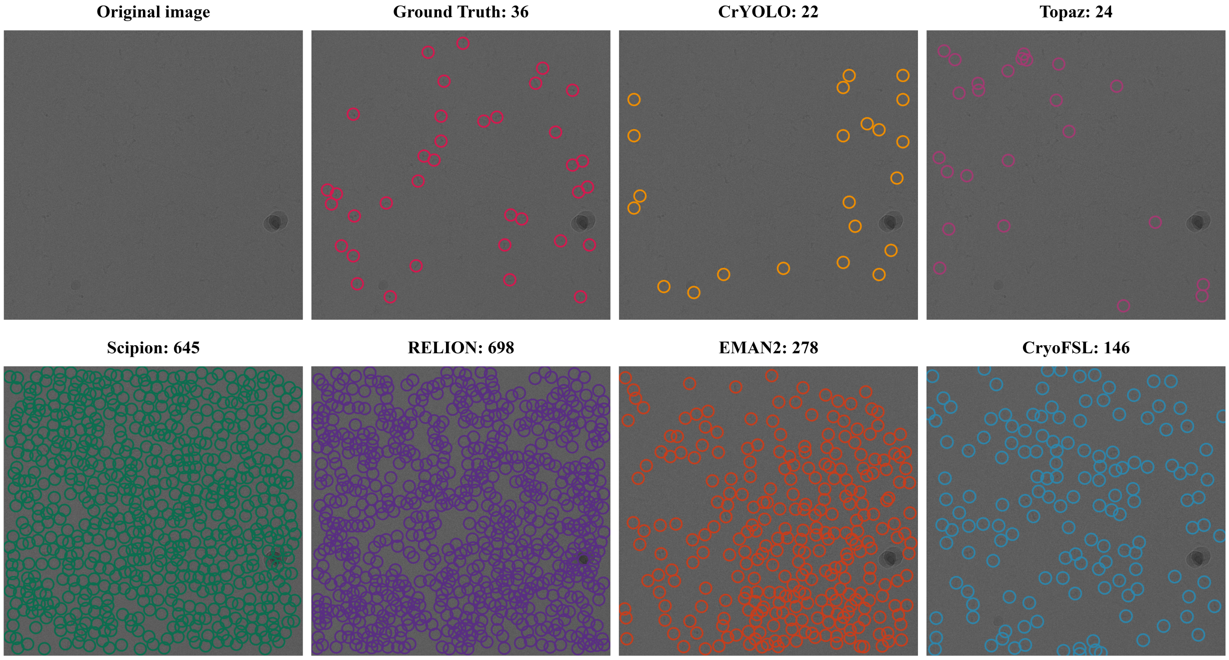


(b)

**Supplementary Figure S6:** Comparative analysis of particle detection across methods on EMPIAR-10345 dataset. (a) Displays multiple sample micrographs from the EMPIAR-10345 dataset, comparing GT annotations (left) with CryoFSL predictions (right), with particle numbers annotated alongside. CryoFSL accurately identifies GT particles and detects additional true positives (TPs) missed during GT curation, reflecting its high recall and robustness, though precision is lower due to these extra valid detections rather than false positives. (b) Detailed comparison of all six particle picking methods on sample micrograph 18jam15a_0309_ali_DW, illustrating distinct failure modes across approaches. CrYOLO and Topaz exhibit systematic under-picking, missing substantial numbers of true particles present in the ground truth. In contrast, template-based methods (Scipion, RELION, and EMAN2) demonstrate over-picking behavior, selecting numerous false positive regions that do not correspond to actual protein particles. The lower precision observed in these traditional methods stems from excessive false positive detections, fundamentally differing from CryoFSL's precision characteristics, which result from identifying legitimate particles missed during manual annotation rather than selecting non-particle regions. This comparison highlights the distinct nature of precision limitations across methods: CryoFSL's comprehensive true positive recovery versus traditional methods' false positive contamination.

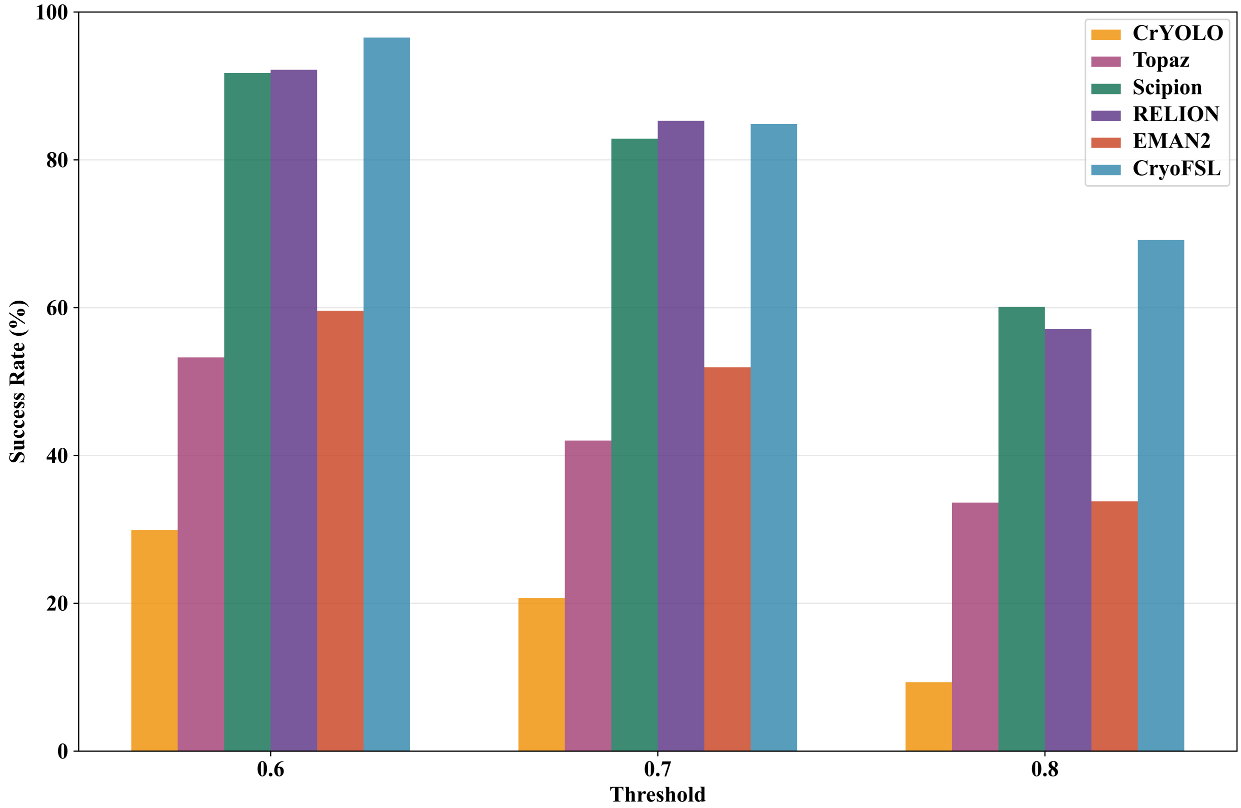


**Supplementary Figure S7:** Bar chart illustrating the recall success rate of different particle picking methods at thresholds 0.6, 0.7, and 0.8. Success rate is defined as the percentage of cases for which a method achieves a recall value equal to or exceeding predefined thresholds. The x-axis indicates the recall threshold values, and the y-axis represents the success rate in %. Each bar corresponds to a method's success rate at a specific threshold, with taller bars indicating a higher proportion of cases meeting or exceeding the recall criterion.

**Threshold-based recall success rate evaluation**

In order to determine stability under varying recall expectations, we computed the recall success rate, defined as the percentage of cases for which a method achieves a recall value equal to or exceeding predefined thresholds (0.6, 0.7, and 0.8). As shown in Supplementary Figure S7, CryoFSL outperforms all baselines across all thresholds, demonstrating not only high recall but also steady performance even under stringent conditions. At the moderate threshold of 0.6, CryoFSL achieved an outstanding 96.6% success rate with minimal variability (±3.9%), indicating near-universal capability to detect the majority of particles. This performance greatly surpasses deep learning methods like Topaz (53.3 ± 34.1%) and CrYOLO (29.9 ± 40.0%), while closely matching template-based methods such as RELION (92.2 ± 13.5%) and Scipion (91.7 ± 17.3%).

At a stringent threshold of 0.8, which represents a high confidence requirement for particle detection, CryoFSL sustains a 69.1% success rate, representing a 9.1% performance advantage over the next-best model (Scipion at 60.1%) and a staggering 7.4-fold improvement over CrYOLO (9.3%). The sharp decline in success rates for conventional methods as the recall threshold increases highlights their inconsistency in achieving recall. The steep variance (e.g., ±36.2% for Topaz and ±38.6% for EMAN2 at 0.8) further exposes the inconsistency in their results, with success highly dependent on the specific proteins or datasets. This clearly illustrates that our approach maintains reliable particle detection even under high-confidence requirements, setting it apart in terms of dependability and practical utility.

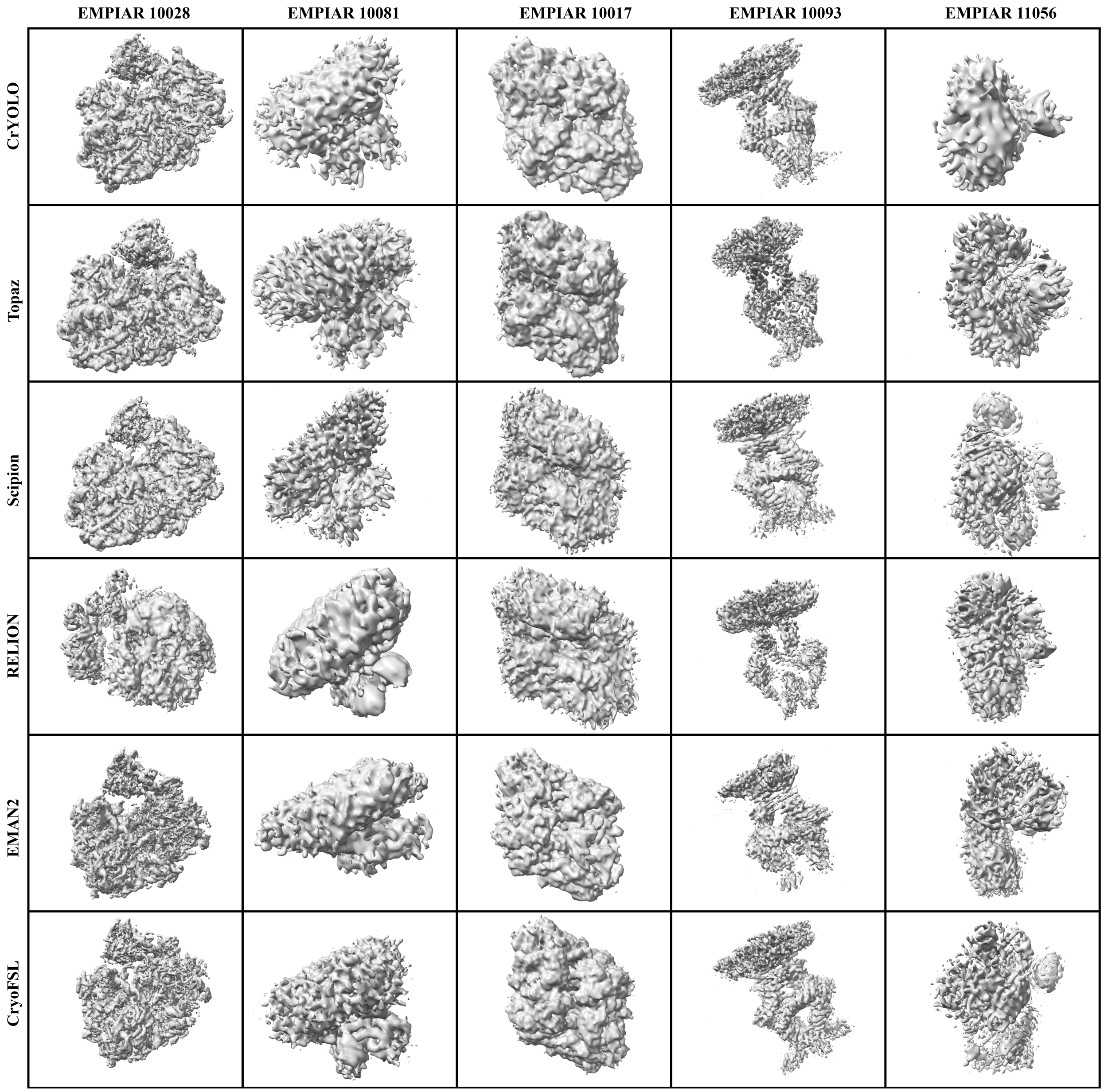


**Supplementary Figure S8:** Comparison results for density maps of particles picked by different methods across multiple EMPIAR IDs (10028, 10081, 10017, 10093, and 11056). CryoFSL outputs high-resolution density maps compared to other methods in most protein types.

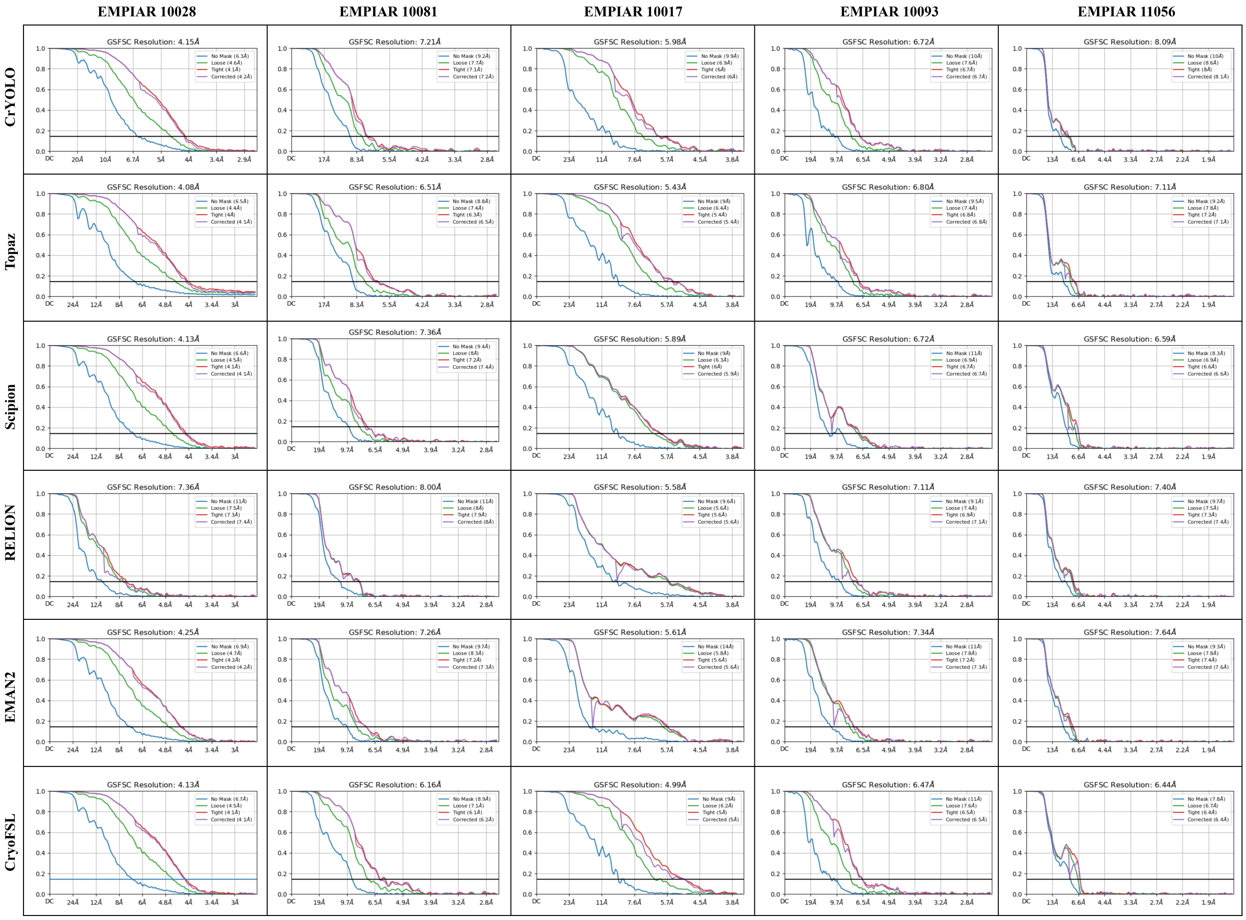


**Supplementary Figure S9:** Comparison results for resolution of density maps of particles picked by different methods across multiple EMPIAR IDs (10028, 10081, 10017, 10093, and 11056). CryoFSL has better resolution than other methods in most protein types.

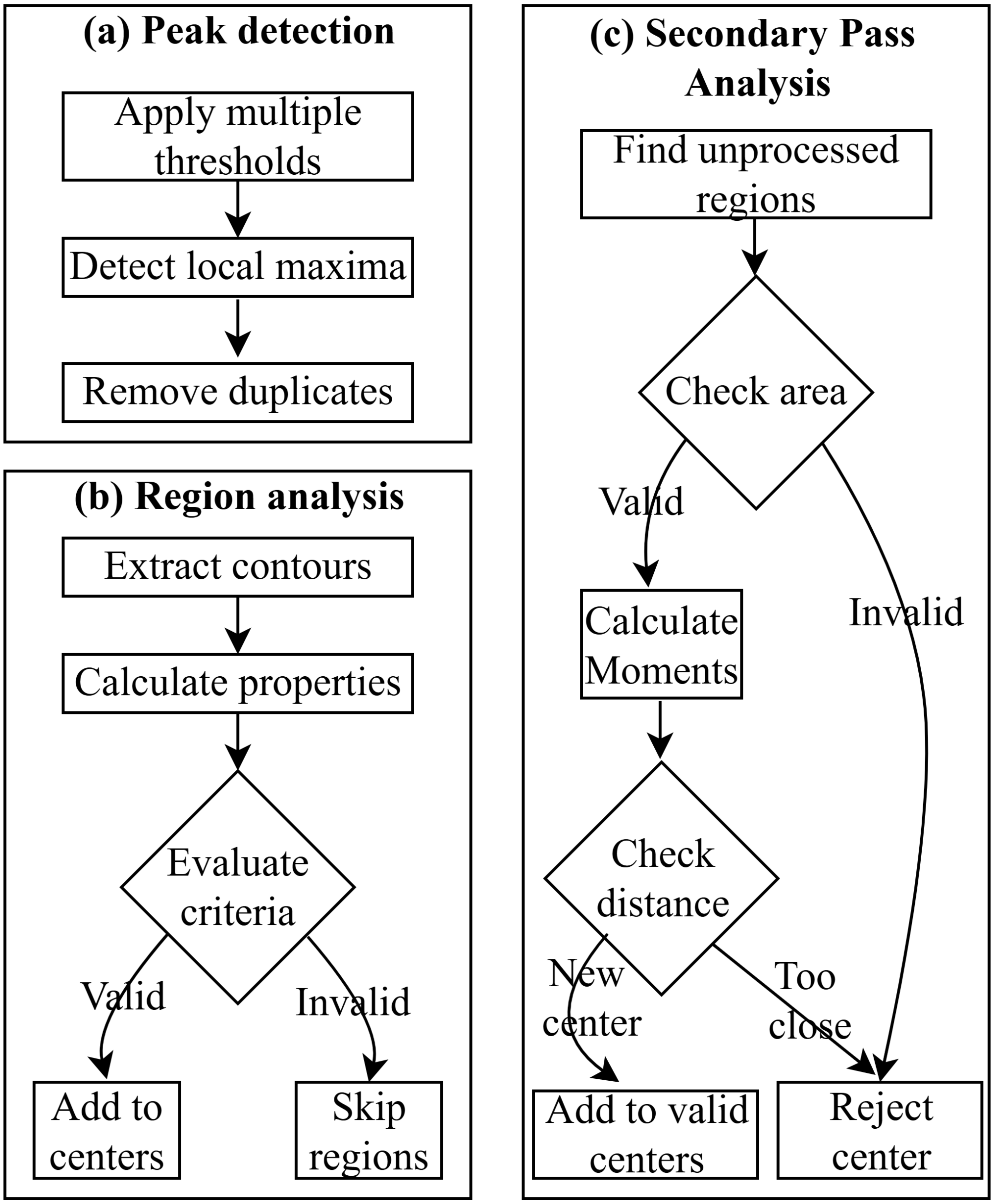


**Supplementary Figure S10:** Flowchart of the enhanced particle center generation algorithm for post-processing in CryoFSL. The pipeline consists of three main stages: (a) Peak detection module that applies multiple distance transform thresholds to detect local maxima and removes duplicate detections, (b) Region analysis module that extracts contours from watershed-segmented regions, calculates geometric properties, and validates regions based on area and circularity criteria, and (c) Secondary pass analysis module that identifies unprocessed regions and applies additional validation based on area and distance constraints to detect previously missed particles. Diamond shapes represent decision points with validation criteria, rectangular boxes indicate processing steps, and arrows show the sequential flow of operations. Valid regions are added to the final particle center list, while invalid regions are rejected or skipped. The complete algorithmic implementation with detailed parameters and mathematical formulations is provided in Supplementary Algorithm S1.

### 1 Supplementary Algorithm S1: Enhanced particle center generation

**Input:** Binary mask Image $B(x, y) \in (0, 1)$, expected particle radius $r$
**Output:** List of particle centers $\{(x_1, y_1), (x_2, y_2), \ldots, (x_n, y_n)\}$

1. Calculate the distance transform:
   $D(x, y) = \min\left\{\sqrt{(x - x')^2 + (y - y')^2} \mid B(x', y') = 0\right\}$
2. Multi-scale peak detection
   (a) Define thresholds $T_k = \alpha_k \cdot r$, where $\alpha_k \in \{0.1, 0.15, 0.2, 0.25\}$
   (b) For each $T_k$:
      i. Detect local maxima: $C_k = \{(x, y) \mid D(x, y) > T_k\}$
      ii. Store coordinates of peaks and remove duplicate coordinates: $C = \bigcup_k C_k$
3. Create markers $M(x, y)$ where each unique center in $C$ is assigned a distinct label.
4. Perform watershed segmentation on $-D(x, y)$ using $M(x, y)$:
   $L(x, y) = i$, if $(x, y)$ belongs to the $i^{\text{th}}$ region.
5. For each region $R_i$ in $L(x, y)$:
   (a) Compute area $A$, perimeter $P$, and circularity Cir.
   (b) Validate regions based on $A \in [0.3\pi r^2, 2.2\pi r^2]$ and Cir $> 0.5$ or $A < 1.2\pi r^2$
   (c) For valid regions, compute centroids using moments:
   $$(c_x, c_y) = \left(\frac{M_{10}}{M_{00}}, \frac{M_{01}}{M_{00}}\right)$$
6. Create mask of unprocessed regions by subtracting processed regions from input mask
7. Detect new centers based on area and distance validation from existing centers
8. Returns the list of valid particle centers
